# Genomic ancestry predicts rapid responses to drought across spatiotemporal scales

**DOI:** 10.1101/2025.10.02.679894

**Authors:** Rozenn M. Pineau, Natalia Bercovich, Loren H. Rieseberg, Julia M. Kreiner

## Abstract

How genetic diversity responds to environmental change across spatiotemporal scales remains poorly understood despite its importance for species persistence in changing landscapes. Agricultural weeds offer ideal models for studying these adaptive dynamics as they rapidly evolve under both the intensive management practices designed to eliminate them and increasingly severe climate challenges such as drought. Here, we combine experimental and herbarium genomic approaches spanning within-generation to century-long timescales to understand how genome-wide variation responds to drought in *Amaranthus tuberculatus*. In this native species, a history of divergent selection between two ancestral lineages followed by secondary contact is thought to have facilitated its invasion into agriculture. A drought survival experiment on accessions from paired agricultural and natural populations across its range revealed substantial phenotypic variation differentiated by habitat, geography, and ancestry. Ancestry mapping revealed 43 independent regions across nearly all chromosomes that confer protective effects under drought, demonstrate particularly rapid allele frequency changes, and exhibit duration-specific selection over the course of the imposed drought. Observation of allele frequencies across the past century reveal evidence for climate-dependent fluctuating selection governing the evolution of drought-associated loci. Selection favors drought alleles during hot/dry years and selects against them in cool/wet years—a pattern more evident in long-term trends than in shorter temporal intervals, suggestive of adaptive lag in rapidly changing environments. By combining short and long-term spatiotemporal data, we demonstrate that fluctuating selection has preserved the polygenic variation underlying population responses to drought, enabling ongoing adaptive responses to contemporary land-use and climate change.

**Significance Statement:** Understanding how species cope with rapid climate and land use change requires studying evolutionary responses across scales. Using *Amaranthus tuberculatus*, a native species turned major agricultural weed, we bridge timescales by pairing a drought experiment with century-spanning herbarium genomics. We show that ancestry structures fitness under drought and has driven agricultural populations to be better drought-adapted. This involves many genes whose allele frequencies fluctuate with climate: drought-protective alleles increase during hot/dry years and decline in cool/wet years. These fluctuations maintain genetic diversity and enable climate tracking, which is imperfect over short timescales. By linking experimental and historical data, we uncover evolutionary dynamics missed by snapshots, improving predictions of species adaptation to environmental change and informing weed management.

## Introduction

Human activities are driving environmental change at unprecedented rates, creating intense and rapidly fluctuating selection pressures that challenge traditional evolutionary theory. The ‘standard genetic model’ assumes selection is weak, gradually depletes variation, and drives slow evolutionary change (1, 2). This view, however, fails to capture the intensity and fluctuating nature of human land-use practices and climate change, which create spatially and temporally variable fitness landscapes across contemporary environments. When selection varies across space and time, fitness trade-offs mediate the balance between global evolutionary forces and local adaptive optima (3, 4). The rate at which populations adapt locally depends on the strength of selection relative to migration (5, 6) and the availability of adaptive genetic variation, with standing variation facilitating rapid responses (7). Understanding the implications of environmental change on adaptation and species persistence therefore requires quantifying how genetic variation responds to selection across relevant spatial and temporal scales.

While short-term responses can be measured across generations, they rarely predict long-term evolutionary dynamics (8–10). Temporal genomic field experiments have demonstrated that nonlinear and fluctuating selection can slow trait evolution while maintaining genetic variation (4, 11–13), and that climate-driven selection operates across seasonal to decadal timescales (13–16). The sequencing of historical herbarium samples spanning centuries of environmental change offer a powerful approach to address these phenomena in plant species (17–19), and has been leveraged to understand the consequences of changing land-use (20) and climate (21, 22). However, we still lack a clear understanding of how short-term adaptive responses under intense selection relate to longer-term evolutionary dynamics across contemporary landscapes (23, 24).

A particularly important adaptive response in plants occurs under drought, which imposes strong selective pressure through prolonged water limitation (25, 26) that varies dramatically in space, time, and intensity (27, 28). This spatial heterogeneity has long been recognized as important for plant ecotype formation (29, 30), with locally adapted types leveraging diverse physiological and biochemical responses ranging from altered phytohormone signaling to resource allocation (31, 32). These mechanisms constitute distinct but potentially co-occurring ecological strategies: drought escape (reproducing before drought), avoidance (maintaining hydration), and tolerance (withstanding dehydration) (28). Given the complexity of strategies and traits involved, it is not surprising that drought adaptation is often polygenic (e.g. (33–35)). This architecture might further be expected to vary across populations due to locally varying optima (36), especially as adaptation to drought frequently imposes costs by reducing growth and reproduction (37, 38). Therefore, the extent and architecture of drought adaptation may differ considerably across environments with varying water availability, competition levels, and life history demands as is typical of contemporary landscapes, where human-mediated disturbance creates a mosaic of selection regimes.

Waterhemp (*Amaranthus tuberculatus*) is a species native to North America which has successfully expanded from wetland habitats into cultivated lands in the second half of the 20th century (39), presenting a powerful system for studying rapid adaptation (40, 41). Two varieties within the species, var. *rudis* and var. *tuberculatus*, were historically isolated from one another by the Mississippi River but have been brought back into secondary contact (40, 42). Var. *rudis* ancestry has been favored during the species’ invasion into cultivated lands (41), consistent with hypotheses that it was preadapted to agricultural habitats (40). Genomic time series from herbarium specimens have revealed how this ancestral standing genetic variation facilitated waterhemp’s rapid response to agricultural intensification (20). However, we do not know what historical selective pressures maintained the genetic variation in var. *rudis* that enabled its success in agricultural environments (43). Given climatic differences across the species range, we hypothesized that higher exposure to drought in var. *rudis’* evolutionary history facilitated waterhemp’s transition from riparian wetlands to drier agricultural landscapes.

Here, we test this hypothesis by characterizing variation in adaptation to drought across *A. tuberculatus*’ native range at both genomic and phenotypic scales. To do so, we first conducted a controlled drought survival experiment on range-wide accessions collected from paired agricultural and natural habitats, allowing us to disentangle the effects of geography, habitat type, and ancestry on drought avoidance. Using whole-genome sequencing of 280 individuals from this experiment, we employed admixture mapping to identify genomic regions underlying drought adaptation and tracked their allele frequency dynamics and phenotypic effects throughout an extended drought treatment. To understand the evolutionary history of these adaptive loci, we analyzed climate-mediated selection patterns using herbarium specimens spanning over a century of environmental change—testing whether genomic regions important for drought survival today show signatures of selection over historically changing climates.

## Results

### Drought avoidance across the landscape varies by geography and environment

To understand how extreme bouts of selection from drought of varying duration impact phenotypic and genomic variation across the landscape, we conducted a survival experiment. This experiment drew from accessions spanning 24 *Amaranthus tuberculatus* populations, from natural and agricultural habitats that were collected pairwise across the Midwestern United States (12 pairs total, **Fig. 1A)** (41). After 3 weeks of growth (the 4-6 leaf stage), we stopped watering plants in the drought but not control treatment (implemented in a randomized full block design). We monitored survival by recording the day to full plant wilt (**Fig. 1, Fig. S1**)–a proxy for survival that could reflect adaptive variation in drought avoidance. In a mixed effects model of an individual’s survival, controlling for block effects (bench, tray position effects) and size at imposed drought, we found that longitude significantly explained individual level variation in drought avoidance (β= −0.13, t = −2.427, *p*-value = 0.016).

**Fig. 1.**
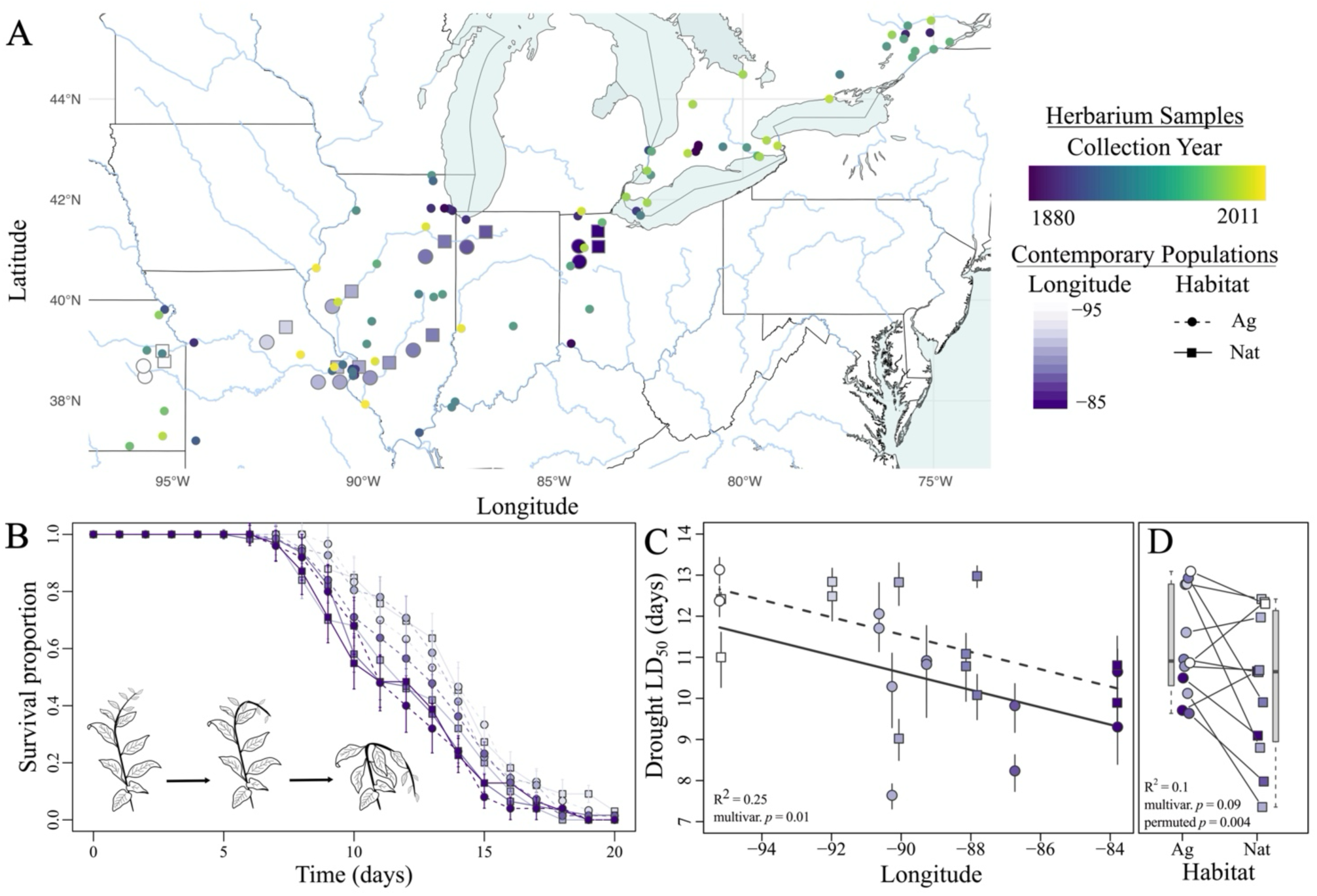
Drought avoidance varies by habitat and across the geographic range of A. *tuberculatus*. **(A)** Paired agricultural and natural populations were collected on a southwestern to northeastern gradient, on both sides of the Mississippi river—spanning the ancestral ranges of *var. rudis* and *var. tuberculatus*. Samples are colored by longitude, while environment is indicated by shape (see legend). Smaller circles denote the sequenced herbarium specimens’ sampling location (from Kreiner et al., 2022), with color representing year of collection. **(B)** Plants from contemporary collections were subjected to a drought selection experiment and their survival over time, based on a proxy of complete turgor pressure loss (see inset), was recorded. Points represent means grouped by longitude and environment for visualization purposes (legend as in A, error bars represent the standard error). **(C)** The population level drought LD_50_ (days until 50% mortality) is higher in eastern compared to western populations, and in **(D)** populations collected from agricultural compared to nearby natural habitats. Partial R^2^ values correspond to a multivariate model testing the effect of habitat and longitude on drought LD50 (**Table S1**).

Given our paired agricultural-natural sampling design, we further probed whether there were consistent differences in drought avoidance among populations depending on land-management—whether human-mediated agricultural or unmanaged natural lands. From population-specific survival curves, we determined the population drought-LD_50_, defined as the number of days of drought exposure required to reach 50% mortality in the population (**Fig. 1B, Fig. S2**). Permuting habitat assignment among populations, we found that populations collected in agricultural habitats were significantly more resilient to drought as compared to those from natural habitats (permuted *p*-value = 0.004; **Fig. 1B-D**, **Fig. S3**). A multivariate regression of population LD_50_ with only environment and longitude supported this directionality (**Table S1**), with the least square mean LD_50_ in agricultural habitats equal to 11.4 days (95% CI = [10.67, 12.13]), as compared to 10.5 days (95% CI = [9.77, 11.23]) in natural habitats (β=-0.93, R^2^ = 0.10, F_1,21_ = 3.14, *p*-value = 0.09). Additionally, it recovered a longitudinal gradient in LD_50_ values, indicating better adaptation to water limitation in our most western populations (12.0 days, 95% CI = [10.45, 13.55]) compared to easternmost populations (10.1 days, 95% CI = [8.80, 11.40]) (**Fig. 1C**, β=-0.21, longitude partial R^2^ = 0.25, F_1,21_ = 8.2, *p*-value = 0.01), consistent with parallel population responses to drought at geographic and environmental scales.

Ecological trade-offs often constrain species’ responses to environmental stress (24). Rapid growth can enable drought escape by completing development before severe water stress occurs, potentially reducing selection for physiological drought avoidance mechanisms. Herbicide resistance mechanisms like EPSPS gene amplification can additionally impose growth costs (44), suggesting potential tradeoffs among growth, drought responses, and herbicide resistance. We investigated these relationships using early growth rate measurements from the survival experiment and genomic estimates of EPSPS copy number. Individuals varied considerably in EPSPS copy number, with 61.5% showing no amplification and 38.5% showing amplification up to 5.51 copies. At the individual level, univariate correlations revealed positive relationships between drought avoidance and both growth rate (β = 0.74, Pearson’s r = 0.33, 95% CI = [0.21, 0.44], *p*-value = 5.5×10^-7^, **Fig. S4 & Table S2**) and herbicide resistance (β = 0.05, Pearson’s r = 0.16, 95% CI = [0.04, 0.27], *p*-value = 0.009, **Fig. S4 & Table S2**). In mixed-effects models of drought avoidance that controlled for experimental block structure (greenhouse bench and tray), and accounted for geographic origin and habitat, only the relationship between drought avoidance and growth rate remained significant (β = 0.77, t = 5.34, *p*-value = 2.4×10^-7^, **Table S3**), while the association with herbicide resistance disappeared (**Fig. S4** and **Table S3**). These patterns contradict expectations for both growth-resistance tradeoffs and for the drought escape-avoidance dichotomy.

Despite these individual-level correlations, population-level comparisons revealed clear habitat-associated structure in trait distributions. Mean trait values in agricultural populations show not only increased drought avoidance as described above, but also higher frequencies of herbicide resistance (**Fig. S5C**; Wilcoxon signed-rank test, *p*-value = 0.0002), and slower early growth rates compared to natural populations (**Fig. S5**; Wilcoxon signed-rank test, *p*-value = 6.80 x 10^-7^). This scale-dependent pattern, where habitat differences in trait means contradict individual-level correlations, suggests that population structure and correlated selection may underlie trait divergence between agricultural and natural populations.

### Ancestry predicts response to drought

To understand how drought shapes genome-wide variation, we resequenced and analyzed whole-genomes from 280 individuals subjected to drought in our survival experiment. We found strong signatures of population structure consistent with previous work (20, 41). The first and second principal components (PC) of an ordination on genome-wide genotypes explained 17% and 6% of the variation among samples, and were strongly predicted by several geographic and environmental variables in multivariate regressions (*PC1 predictors*: latitude F_3, 276_ = −8.94, *p*-value < 2 x 10^-16^, partial R^2^ = 0.08; longitude F_3, 276_ = −9.83, *p*-value < 2 x 10^-16^, partial R^2^ = 0.12; and environment F_3, 276_ = −4.37, *p*-value = 1.8 x 10^-5^, partial R^2^ = 0.02; *PC2 predictors*: environment F_3, 276_ = −4.12, *p*-value = 5 x 10^-5^, partial R^2^ = 0.06; **Fig. 2A, Fig. S6**). An ADMIXTURE analysis (45) with two ancestral populations (K=2) was the best supported aside from K=1 (**Fig. S6 B&C),** and was almost perfectly collinear with PC1 (Pearson’s r = 0.992, CI = [0.990, 0.994], *p*-value < 2.2e-16, **Fig. 2B** & **Fig. S7**). While additional population substructure was present at higher values of K and across higher PCs (**Fig. S6)**, the primary axis of structure is consistent with two ancestral groups, *A. tuberculatus* var. *rudis* and var. *tuberculatus*, to the southwest and northeast, respectively (**Fig. 2A**).

**Fig. 2.**
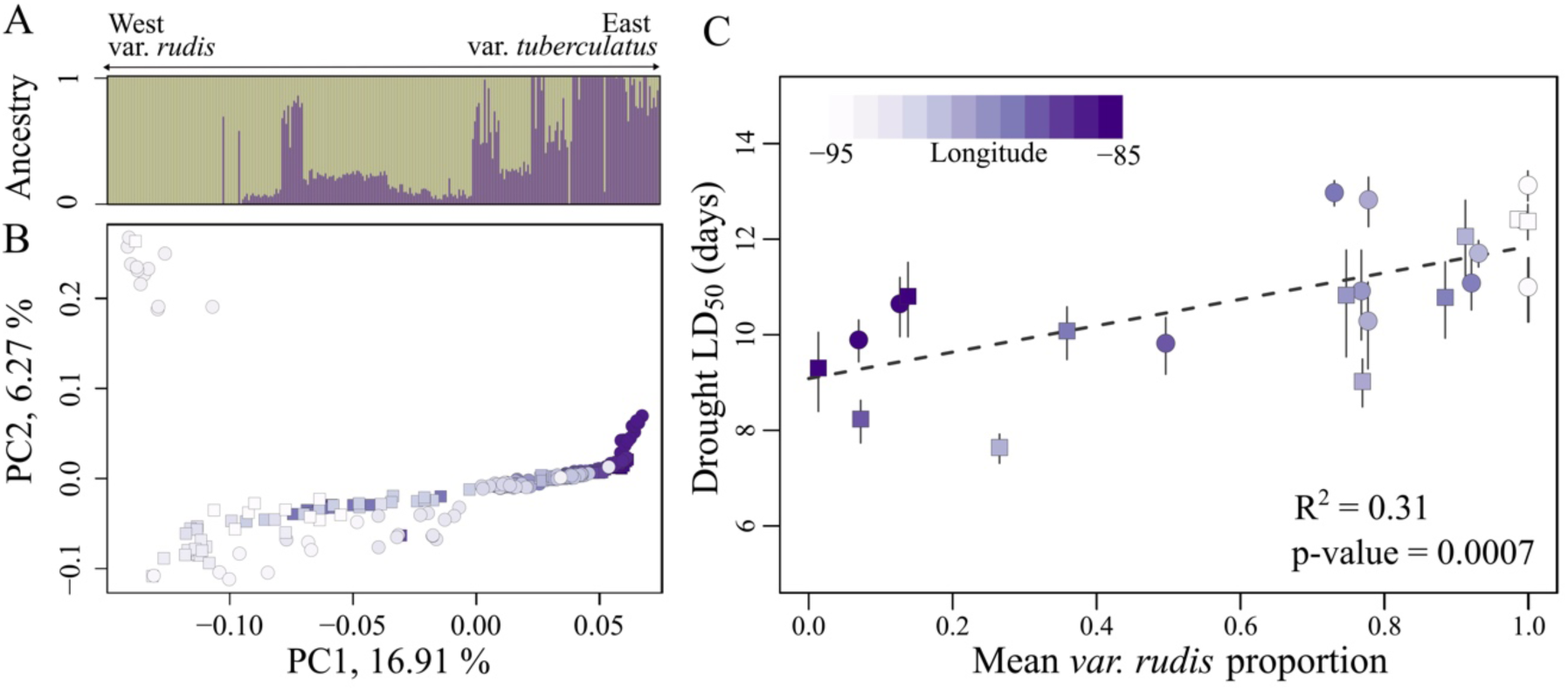
Genome-wide ancestry predicts the response to drought. **(A)** Model based global ancestry inference (ADMIXTURE) demonstrates that our experiment spans a range of individual-level ancestry composition, with a gradient of ancestry going from pure var. *rudis* in the west to pure var. *tuberculatus* in the east. Samples are ordered by longitude. **(B)** Population structure as inferred by principal component (PC) analysis, with the first PC showing clear structure by longitude, such as we observe for admixture. **(C)** The mean proportion of var. *rudis* ancestry in a population significantly predicts population drought LD_50_. Samples are colored by longitude in B&C, while squares and circles represent natural and agricultural populations respectively.

We hypothesized that geographic and habitat-based patterns in drought avoidance may be explained by variation in ancestry composition. Indeed, we found that the mean *var. rudis* ancestry proportion in a population is positively associated with drought LD_50_ in a multivariate regression regardless of whether we use an ADMIXTURE or PC based ancestry metric (**Table S4)**. This occurs both through a main effect and through ancestry’s interaction with longitude (**Fig. 2C**, ancestry: F_1,16_= 5.96, *p*-value = 0.03, partial R^2^ = 0.31; longitude x ancestry: F_1,16_ = 7.15, *p*-value = 0.017). This relationship was also detected at the individual level when testing the effect of ancestry on the day to full wilt in a multivariate model (**Fig. S8,** ancestry: F_1,276_= 36.99, *p*-value = 0.05, ancestry partial R^2^ = 0.033; longitude x ancestry: F_1,276_ = 43.07, *p*-value = 0.03). The longitude by ancestry interaction reveals that longitude predicts variation in drought LD_50_ among populations when var. *rudis* ancestry is rare or absent, but has no effect when var. *rudis* ancestry dominates (**Fig. S9)**. Near the center of the range where ancestry varies considerably (−88° to −91°), a shift from pure var. *tuberculatus* to pure var. *rudis* results in 1.7-2.3 fold increase in LD_50_. These results support that divergent evolutionary histories between lineages have produced substantial heritable variation in drought avoidance, providing a foundation for mapping the genomic architecture of drought.

### Resolving the genetic architecture of drought avoidance

Given the strong association between var. *rudis* ancestry and drought avoidance in our system, we expected that a genome-wide association study controlling for ancestry would result in a high false negative rate (**Fig. S10**). We therefore leveraged an admixture mapping approach to identify tracts of ancestry across the genome underlying adaptation to drought. We performed local ancestry inference on our 280 resequenced samples using ancestry_hmm (46) (see *Methods*), leveraging 786,262 ancestry informative variants (∼1.2 per kb) to call fine-scale ancestry across the genome (**Fig. 3A**). Under a two-pulse admixture model, our data is consistent with a first pulse of admixture 10,000 generations ago that introduced 0.42 proportion of var. *rudis* ancestry into var. *tuberculatus*, and a second pulse 7.4 generations ago which introduced 0.24 proportion of var. *rudis* ancestry. This fine-scale inference was highly concordant with estimates of genome-wide level ancestry inference, reinforcing the considerable variation in ancestry across individuals (**Fig. S11**).

**Fig. 3.**
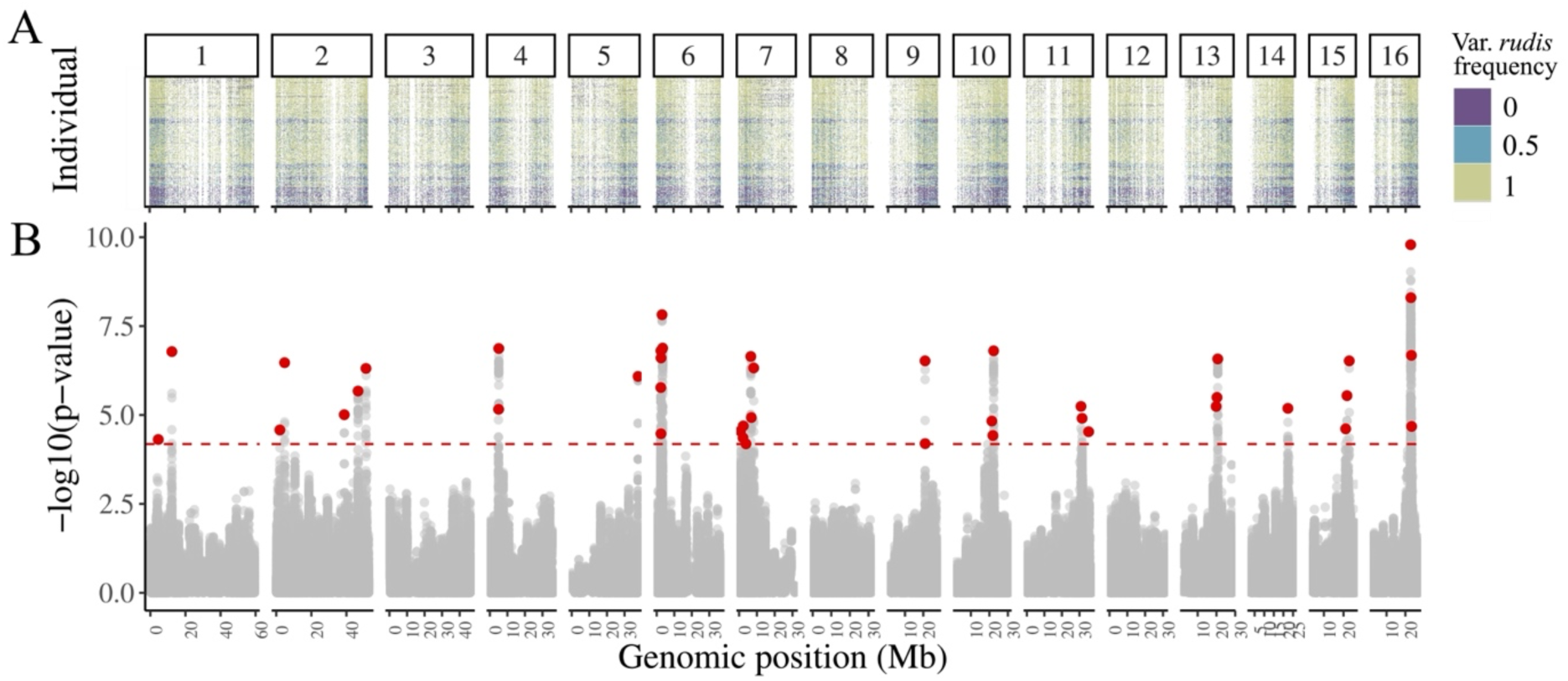
Admixture mapping reveals the polygenic architecture underlying drought avoidance. **(A)** Fine-scale ancestry calls for 786,261 loci across the genome resulting from a two-pulse admixture model, with individuals ordered by longitude along the y axis (bottom to top, western to eastern). **(B)** Admixture mapping of survival under drought reveals 893 significantly associated loci after FDR and genomic inflation factor correction (q <0.05, λ = 2.35, lower dashed horizontal line), mapping to 43 independent loci (focal loci indicated in red dots, based on LD clumping of sites with r^2^ > 0.50 within a 100 kb region).

We identified 893 significant associations between local ancestry and drought avoidance, representing 43 independent loci after LD clumping, FDR, and genomic inflation factor correction (q < 0.05, λ = 2.35, **Fig. S12**). These were ascertained while controlling for population structure using the kinship matrix (see methods; **Fig. 3A**), which yielded a SNP heritability estimate of 10.6% (SE = 1.0%). Given the relatively fast decay of linkage disequilibrium in waterhemp (b = 1.1×10−4, **Fig. S13**) and relatively short half-decay distance (6,539 bp), it is notable that the size of drought associated loci are particularly large, ranging from 6.3 kb to 196.4 kb (mean = 77 kb), spanning 1-25 genes (mean = 6.4 genes) and 9-776 linked drought-associated SNPs (mean = 117.5 SNPs, **Fig. S13**). These drought loci are found across 13/16 chromosomes and many fall within genes previously identified to have drought-relevant functions, including *APSR1* (β = 1.30 & p-value of lead SNP = 1 x 10^-7^) (47), *PER17* (β = 1.24 & p-value of lead SNP = 1 x 10^-6^) (48) and *RKD5* (β = 1.27 & p-value of lead SNP = 7 x10^-7^) (49), *TDC2* (β = 1.14 & p-value of lead SNP = 2 x 10^-6^) (50), and *DDB1* (β = 1.40 & p-value of lead SNP = 8 x 10^-9^) (51–53) (see full list in **Table S5**). *DDB1* is a strong candidate for a particularly large effect locus, as not only does it exhibit the largest effect size and most significant association, but also 9 distinct non-synonymous mutations at drought-associated SNPs. GO biological processes for these drought-associated loci include functions such as mRNA processing (cleavage and splicing), plant growth (gravitropism, photomorphogenesis, root and leaf development), protein synthesis (tRNA), and ion metabolism (ion binding and transport) (**Table S6**). These results are consistent with a complex genetic basis of drought avoidance.

We next tested whether drought associated alleles were enriched in their differentiation among natural and agricultural habitats. Leveraging our paired sampling design, we implemented a Cochran-Mantel-Haenszel (CMH) scan to identify genomic regions involved in agricultural adaptation (as in (20)), identifying 94,008 SNPs spanning 44,692 genes (∼42,831 SNPs after clumping) across the genome surpassing FDR correction (α = 0.01, see *Methods*, **Fig. S14 & Fig. S15**). Intersecting these putatively agriculturally-adaptive alleles with our drought-associated loci, we find that ∼3.4% (30/893) of drought-associated loci are amongst the most divergently selected among natural and agricultural habitats (i.e. loci in the genes *HAT14*, *RKD5*, *TBR*, *TDC2*). This overlap, however, did not exceed null expectations based on the observed distribution of ancestry tract lengths, suggesting drought alleles were not consistently among the most differentiated agricultural-natural loci genome-wide (**Fig. S16**). While drought tolerance does not appear to be a universal driver of agricultural adaptation, habitat significantly predicted drought allele frequencies after controlling for genome-wide ancestry (main effect: F_1,271_ = 4.34, *p*-value = 0.038; habitat × latitude: F_1,271_= 4.45, *p*-value = 0.036; habitat × longitude: F_1,271_ = 4.61, *p*-value = 0.033). Agricultural populations showed elevated drought allele frequencies relative to natural populations, with the strongest enrichment in western (Δp = 0.062) and northern (Δp = 0.088) regions (**Fig. S16**). This suggests that agricultural environments favor these drought adapted alleles, but with selection strength varying geographically.

### The tempo of selective responses across an extreme drought event

Our drought survival experiment provides insight into the rate and scale of genomic response to selection imposed by extreme climate events. By tracking the 43 independent drought-associated loci over the progression of the imposed drought, we capture the selected allele frequency trajectories contributing to drought adaptation in *A. tuberculatus*. We find that the frequency of var. *rudis* haplotypes at all 43 drought-associated regions increase over the course of the selection experiment (**Fig. 4A&B**), providing further evidence for the protective effect of var. *rudis* ancestry under water limitation. This ancestry-specific response emerged despite a high background frequency of var. *rudis* (mean var. *rudis* frequency at the start of the experiment ∼0.73) in our experimental design. To further distinguish selection at these sites from shifts in the genome-wide background, we contrasted the mean change in the frequency of these 43 drought-adapted loci against permutations of 43 randomly selected sites with a matching starting allele frequency distribution (see *Methods*). We found an excess of change in drought-associated loci relative to these frequency-matched distributions (Z score = 1.47, *p*-value = 0.05; **Fig. 4B**), consistent with particularly strong selection imposed on these alleles in our experiment.

**Fig. 4.**
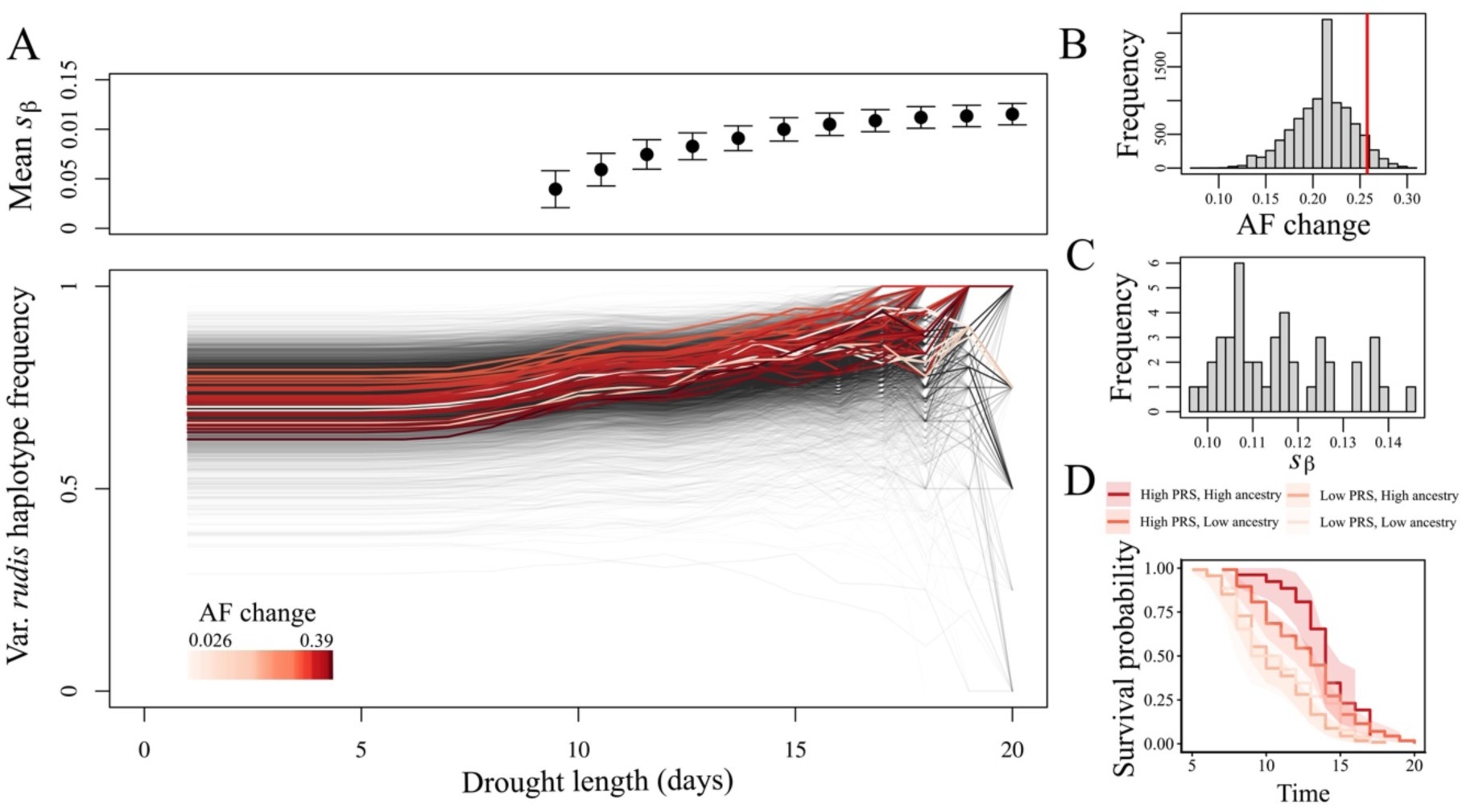
The genomic impact and tempo of drought adaptation across an extreme drought event. **(A, lower panel)** The frequency of var. *rudis* haplotypes over the course of the drought experiment, with the frequencies at day 0 representative of the starting distribution of ancestry among all contemporary samples. Red lines denote the 43 focal loci (i.e. after performing the LD-based clumping algorithm), colored by allele frequency change (Δp̄ = AF_t=20_-AF_t=0_). Black lines reflect 10,000 randomly selected sites to illustrate genome-wide trajectories. **(A, upper panel)** The mean survival-based selection coefficient (*s*_β_) for the 43 drought loci initially increases with drought duration but shows a threshold effect. **(B)** The mean allele frequency change for the 43 drought-adapted alleles (Δp̄ = 0.26, vertical red line) is significantly elevated compared to the genome-wide null expectation (gray distribution; *p*-value = 0.05). **(C)** Within-generation survival-based selection coefficients (*s*_β_) for each drought-associated locus over the course of the entire drought experiment (*s̄*_β_ = 0.12, SD = 0.012). **(D)** Kaplan–Meier curves stratified by PGS (high/low relative to median) and genome-wide ancestry (high/low relative to median).

These time-series data revealed the strength of fitness advantages conferred by drought-alleles in our experiment. The within-generation survival-based selection coefficients (*s*_β_) calculated through logistic regression of allele frequencies through time averaged 0.12 [SE = 0.012] and ranged from 0.097 to 0.14 (see *Methods*), indicating strong selection favoring drought-avoidance alleles during the experimental drought event (**Fig. 4C**). If we consider how the length of such drought events mediates these responses, we observe a threshold response—where *s*_β_ intensified significantly between days 10 and 15 (twofold increase), before stabilizing after day 15 (**Fig. 4A**, top panel). This pattern suggests that drought adaptation involves a non-linear response to water limitation, with selection pressure accelerating dramatically between days 10-15 as plants exceed physiological avoidance thresholds, followed by a stabilization phase after day 15 when surviving individuals become enriched for drought-adaptive alleles.

To partition survival effects attributable to the joint contribution of significant drought-associated loci versus genome-wide ancestry (capturing variants below detection thresholds), we implemented Cox proportional hazards tests. To address potential non-independence among the 43 drought-associated loci and avoid overfitting, we used lasso Cox regression to simultaneously fit all candidate loci and obtain regularized effect sizes. Lasso selected 12 independent loci from the 43 candidates with regularized effect sizes ranging from 0.036 to 0.506. We then calculated polygenic scores using these lasso-regularized effects (PGS = Σ[βᵢ × Aᵢ] (54), where βᵢ represents the lasso-regularized effect size for locus i and Aᵢ represents the var. *rudis* ancestry dosage at that locus), which varied from 0.16 to 4.22. In a Cox proportional hazards model including both PGS and genome-wide ancestry, the PGS was significantly associated with increased survival (hazard ratio = 0.32 per unit, *p* = 5.6×10⁻^18^), while genome-wide ancestry explained additional variance (hazard ratio = 0.11 per unit, *p*-value = 1.3×10⁻^8^) (**Fig. 4D)**. Standardizing by their respective scales, each 10% increase in the PGS was associated with a reduced mortality risk of 37%, while each 10% increase in genome-wide ancestry was associated with a reduced mortality risk of 20% (**Fig. 4D**). As effect sizes were estimated and tested in the same experimental population, these estimates likely represent upper bounds on predictive effects. Nonetheless, the retention of genome-wide ancestry as a significant predictor indicates that additional drought-tolerance variants segregate beyond those captured by the 12 independently-acting loci, demonstrating a complex genetic architecture combining detectable moderate-to-large effect loci on a polygenic background.

### Century-long dynamics of modern drought-loci in response to climate

To study the dynamics of polygenic drought adaptation over the last century, we leveraged herbarium specimens spanning 120 years of changing climate. Leveraging a collection of previously sequenced herbarium specimens (20), we subsampled 87 collected between 1880 to 2011 (**Fig. 1A**) that spans NOAA climate records at fine spatiotemporal scales. After inferring ancestry tracts along the genome in these historical samples, we identified 680 regions that tagged drought-associated loci as identified from admixture mapping, resulting in 33 independent loci after LD-clumping. We analyzed change at these loci alongside NOAA records of maximum temperature and total precipitation in waterhemp’s summer growing season dating back to 1880 (see *methods*, **Fig. S17**).

To investigate whether loci underlying drought-avoidance show evidence of adaptive tracking, we first tested how changes in allele frequency correlated with climate change over short timescales. For this approach, we analyzed pairs of individuals that were geographically close (<100km, **Fig. 5A**) but spanned distinct temporal intervals (ranging from 1 to 111 years apart, **Fig. 5B**), calculating Spearman correlations (ρ) between change in max growing season temperature or change in total summer precipitation and allele frequency change across the 31 drought loci with sufficient data. A Wilcoxon signed-rank test revealed that the distribution of ρ was significantly shifted toward positive values for temperature (*p*-value = 0.003, **Fig. 5C**), but not precipitation, indicating systematic increases in drought-protective alleles under warming conditions and decreases under cooling conditions. Permutation tests (randomizing temperature amongst genotypes) showed that a result at least this extreme occurred 7% of the time under null expectations (permuted *p*-value = 0.07, 95% CI = [0.05, 0.09]). Other correlation metrics showed even more marginal significance, with 3/31 loci showing significant positive correlations compared to the ∼0.73 expected by chance (permuted *p*-value = 0.07, 95% CI = [0.06, 0.09]) and a median correlation coefficient of 0.16 across all loci (permuted *p*-value = 0.11, 95% CI = [0.09, 0.13]). Together, these findings suggest drought alleles show a detectable but weak signal adaptive tracking of temperature over short timescales, suggesting that they are constrained in the extent that they can increase and decrease as temperature varies from season to season.

**Fig. 5.**
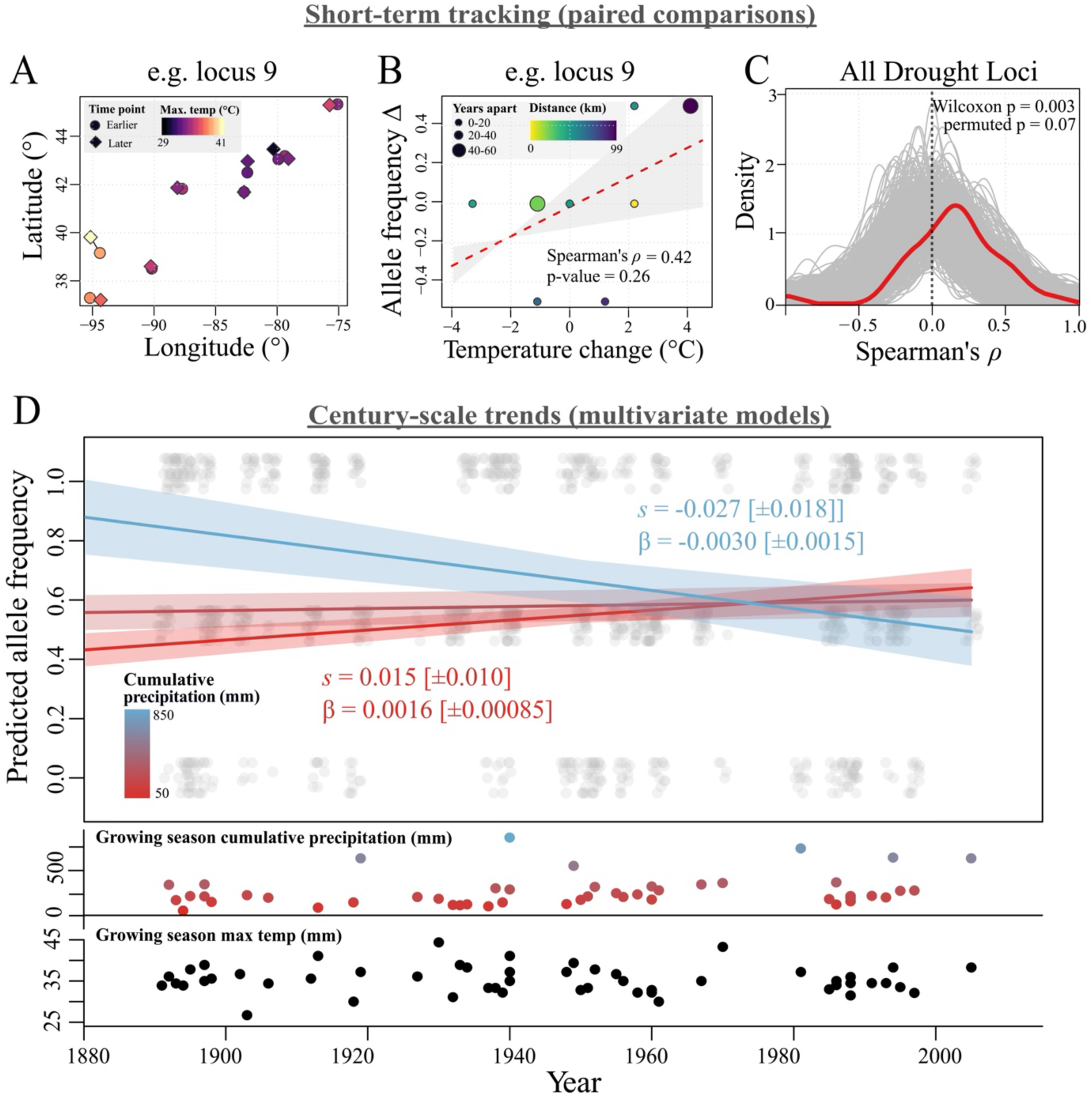
Climate-tracking of drought-adapted alleles across timescales. **(A-C)** Evidence of climate tracking based on correlated shifts in the frequency of drought alleles and maximum temperature (°C) in the growing season between spatially-paired herbarium samples. **(A)** Samples are paired by minimizing distance (km) and proportion of missing data per locus, as exemplified here with drought locus 9. **(B)** The Spearman’s correlation coefficient between temperature change (°C) and allele frequency change is calculated for each set of pairs at each locus, again, exemplified here with drought locus 9. **(C)** The resulting distribution of Spearman’s correlation coefficient observed for 33 herbarium-matched drought loci (red) as compared to a null distribution from permutations randomizing climate with respect to sample (maintaining genotype and temporal structure). **(D)** The collective trajectory of drought loci (predicted allele frequency through time) is precipitation dependent—with evidence of positive selection in dry years and negative selection in wet years. Linear (β) and logistic (*s*) estimates of homozygous fitness advantage from multivariate models including year, year × temperature, year × precipitation, latitude, longitude, habitat, and locus effects. Growing season cumulative precipitation (mm) and growing season maximum temperature (°C) from NOAA corresponding to each spatiotemporal sample is illustrated below.

To gain a comprehensive understanding of the spatiotemporal dynamics of drought adaptation over longer timescales—on average across the last century—we modelled how the collective change in frequency of drought alleles through time and space depends on climate. To do so, we implemented logistic and linear multivariate models of drought allele frequencies that included year, year by maximum growing season temperature, year by total precipitation in the growing season, latitude, longitude, and locus-specific effects. From these models, we extracted selection coefficients (s) and linear regression slopes (β) that are scaled to represent the fitness advantage of homozygous drought allele carriers relative to homozygous reference allele carriers (see *Methods*). The linear analysis revealed significant effects of several predictors (**Table S7**) including the interaction between growing season max temperature and year (β_AA_ = 7.9 x 10^-4^, F_1,1088_ = 8.70, *p*-value = 0.003), indicating that the linear rate of annual allele frequency change is modulated by the temperature experienced in each specific year (**Fig. S18, Table S7**). Based on the logistic model, this temperature effect corresponds to an *s_AA_* of 0.028 [SE = 0.016] for the highest growing season max. temperature regime (45°C) and *s_AA_* = −0.040 [SE = 0.016] for the lowest growing season max. temperature regime (25°C) over the last century (temperature x year effect; X^2^ = 4.71, *p*-value = 0.03).

However, this temperature-driven pattern changed when we considered collection context. After accounting for whether herbarium accessions were found in natural or agricultural settings—which covaries with both temperature and precipitation conditions (**Fig. S19**)—precipitation rather than temperature became the strongest predictor of the linear rate of drought-related allele frequency change through time (β_AA_ = −1.16 x10^-6^, F_1,1002_ = 4.35, *p*-value = 0.037, **Fig. 5D, Table S7).** Furthermore, the temporal response was marginally habitat-dependent (F_2,1002_ = 2.98, *p*-value = 0.05), with agricultural populations showing both faster allele frequency change through time and higher baseline frequencies than populations in natural habitats (F_2,1002_ = 2.97, *p*-value = 0.052). A logistic model recovered weaker signatures of climate-dependent allele frequency change once accounting for collection habitat (growing season cumulative precipitation × year: χ² = 2.34, *p*-value = 0.13), but results were consistent in directionality, with *s_AA_* = +0.015 [SE = 0.010] and *s_AA_* = −0.027 [SE = 0.018] for 50 mm and 850 mm cumulative growing season precipitation regimes, respectively **(Fig 5D)**. Together, these results suggest that human-mediated selection in agricultural environments modulates climate-driven evolutionary responses to drought over century scales.

Beyond these temporal dynamics, drought allele frequencies also showed consistent spatial structuring across climate gradients in herbarium samples. The linear model controlling for habitat revealed that drought allele frequencies differed significantly by precipitation (F_1,1002_ = 4.49, *p*-value = 0.034), longitude (F_1,1002_ = 8.12, p = 0.0045), and latitude (F_1,1002_ = 30.12, *p*-value = 5.13×10^-8^), with among-locus variation also detected (F_32,1002_ = 1.47, *p*-value = 0.045). Under this model, drought allele frequencies are predicted to increase by 6.4 % per degree of latitude. The logistic model yielded qualitatively similar results for the strongest spatial predictors (**Table S7**). This latitudinal gradient is consistent with patterns in our contemporary samples, where northern populations—particularly those in agricultural habitats—exhibit elevated frequencies of drought avoidance alleles. Overall, these spatial patterns complement our temporal findings, together indicating that drought avoidance loci experience fluctuating selection that tracks environmental variation. Selection strength varies predictably across both climatic gradients (latitude, precipitation) and land-use contexts, with agricultural populations showing both stronger responses and higher baseline frequencies of drought alleles.

## Discussion

As global warming intensifies both the severity and frequency of drought events (55), understanding how these changes affect genome-wide diversity within and across populations becomes increasingly critical for conserving and managing biodiversity. To address this challenge, we investigated the genetic architecture and evolutionary dynamics of drought adaptation across the range of a native plant turned agricultural weed, *Amaranthus tuberculatus.* We discovered that variation in drought avoidance correlates with habitat type, geographic location, and ancestry whereby plants from southwestern populations, agricultural habitats, and of var. *rudis* ancestry survive longer under water stress. By leveraging this ancestry-structured phenotypic variation, we identify a polygenic basis for drought adaptation, with drought-related loci distributed across nearly all chromosomes and exhibiting signatures of strong directional selection under extreme seasonal drought events. While this pattern suggests that substantial genome-wide diversity should be depleted under intensifying drought regimes, herbarium genomic analysis of these drought-adapted loci over the past 130 years reveals a signature of fluctuating selection. Collectively, drought-responsive loci exhibit a climate-dependent pattern of selection over time—selected for during hot/dry years but against during wet/cool years—with individual loci varying in their degree of climate-tracking. This dynamic deployment of standing variation suggests that landscape-level variability in anthropogenic selection pressures, including both agricultural practices and drought intensification, can maintain adaptive diversity across spatiotemporal scales.

When environments change rapidly, polygenic responses are expected to draw substantially on standing genetic variation, which facilitates adaptation by enabling more substantial phenotypic shifts than relying on new mutations alone (56–60). While population structure can confound the detection of polygenic architectures (33, 61), here we leverage a strong ancestry-specific response—explaining 33% of the variation in population-level drought avoidance—to map this standing variation and understand the processes that maintain it. Our inference of ancestral haplotypes revealed that ongoing admixture and recombination have broken down large ancestry blocks into fine-scale segments, enabling precise mapping. Admixture mapping identified a polygenic basis for drought adaptation, involving 43 loci across 13 chromosomes (**Fig. 3**), with polygenic scores further revealing substantial variation in the architecture of drought adaptation among individuals. We expect that this variation has been shaped by both the demographic processes that influence the spread of standing variation across landscapes, such as the recent northwestern expansion of var. *rudis* (20, 40), and the adaptive processes that influence their frequency through space and time.

To understand how rapidly variation at these alleles could be lost to selection, we measured the strength of selection acting on these loci during an experimentally-imposed seasonal drought event. During a lethal drought lasting ∼3 weeks, we detect strong directional selection across 43 loci; these regions increase more rapidly than other frequency matched ancestral haplotypes across the genome, at a rate (*s*_β_) of about 12% per day on average (**Fig. 4**). This estimate is akin to a standard selection coefficient (*s)* for the homozygous alternate genotype, but differs in that it reflects the within-generation as opposed to across-generation fitness advantage. We also note that these coefficients may overestimate a true survival-based selection, to the extent that annual plants experiencing prolonged drought can recover from full turgor loss (our experimental proxy for death) with subsequent rainfall. Nonetheless, we expect that these relatively rapid evolutionary responses are enabled by the remarkably large census and estimated contemporary effective population sizes in waterhemp – a common feature of successful agricultural weed species (62, 63). While our observed rates suggest that the hyper-diverse waterhemp can mount the evolutionary responses needed for climate tracking, the critical question is whether such selection strength can be sustained across complex landscapes over decadal timescales.

To quantify these evolutionary dynamics, we examined drought adaptation across spatiotemporal scales using herbarium and past climate records. At a century-long scale, drought allele frequency dynamics reflect an *s* ∼ +0.028/+0.015 in warmest/driest years and - 0.04/-0.027 in coolest/wettest years. The magnitude of the selection coefficients we observe over these timescales are comparable to those estimated as necessary for European *A. thaliana* populations to track predicted 2070 climate conditions (0.01 < s < 0.03; (32)), suggesting waterhemp populations have sufficient adaptive potential to respond to ongoing climate change. Furthermore, these climate-dependent selection coefficients revealed that temporal fluctuating selection has shaped the dynamics of drought adaptation—with costs in wet years exceeding benefits in dry years. Experimentally, we find no evidence of a growth-avoidance tradeoff, highlighting the challenge of detecting fitness costs that may be subtle, environment-specific, or operate through pathways beyond simple growth-survival tradeoffs (66–68). However, costs outweighing benefits for a segregating defense trait have also been described in *Datura wrightii*, where the costs of trichome production outweighed its benefits even in the presence of herbivores (64). In sunflowers, direct selection on rapid growth under benign conditions leads to a loss of drought tolerance (37, 65). Therefore, while temporal fluctuations in selection may be a key mechanism maintaining variation at adaptive loci across species, this mechanism likely works in concert with other factors that impact the efficacy of the response to selection.

Allele frequency trajectories over the past century are influenced by multiple ecological and evolutionary processes that vary across scales (10). Because drought allele frequencies vary predictably with latitude and longitude, suggesting local adaptation to regional climate, the extensive human-mediated long-distance dispersal in waterhemp (66) likely redistributes adaptive variation across environments where it may be locally maladaptive. Buffering effects may also play an important role in the trajectory of these alleles (67). In particular, substantial seed banks and extended dormancy that allow waterhemp seeds to remain viable up to 15 years should slow the traverse of alleles towards a changing optimum (42, 68). Indeed, when examined over shorter timescales using spatially-matched herbarium specimens, drought alleles showed weaker adaptive tracking compared to century-scale trends (**Fig. 5**). This scale-dependent pattern may reflect these buffering processes, though residual spatial variation between paired sites could also contribute. Furthermore, stochastic demographic processes such as boom-and-bust population dynamics can influence allele frequency trajectories in tandem with climate-driven selection (69). The order-of-magnitude population expansion over the last century in waterhemp (63) may have enhanced the effectiveness of selection over more recent timescales, though we have not explicitly modeled these demographic effects. Future work will continue to disentangle the ecological, temporal, and spatial determinants of adaptive allele frequency change, which together have so far been sufficient for rapid responses to changing environments in waterhemp (1, 70).

The scale of the genomic response to drought adaptation likely reflects the complexity of underlying ecological and physiological mechanisms involved. Drought adaptation typically involves escape, avoidance, and tolerance (28, 71) strategies. These can be under the control of few large effect loci, like in drought-escaping *Panicum hallii* (*72*), or a suite of complex traits and accompanying polygenic variation may underlie them in systems such as *Arabidopsis* and *Fragaria* (33, 73, 74). In our experiment, we focus predominantly on avoidance, where several loci with the strongest associations map to DDB1, a protein involved in the abscisic acid hormone signaling pathway in response to drought stress (51), and within *ASPR1*, involved in root growth under nutrient limitation (75). These genes are strong drought avoidance candidates that merit functional validation in waterhemp as transformation protocols improve (e.g. (76)). However, our results suggest a complex mixed-effect architecture—10’s of additional loci show detectable associations (some more broadly involved in the abiotic stress response: *RKD* (49)*, PER17* (48), *TCD2* (*50*)) alongside genome-wide ancestry retaining predictive power—potentially implicating the involvement of other strategies. We were limited in our ability to assess drought escape due to the early initiation of our drought treatment, which killed nearly all plants before flowering. Additionally, our experimental design did not include volumetric water content measurements, preventing us from distinguishing whether survival differences primarily reflect water use efficiency, physiological drought avoidance, or developmental escape strategies. However, we found that more drought-avoidant plants grew faster, and associated variants occasionally map to genes with functions in positive regulation of photomorphogenesis (*TCP2* (77); **Table S6**) and growth (e.g. HAT14 (78))), potentially consistent with drought escape. Future work should incorporate detailed physiological measurements alongside genomic approaches to resolve the mechanistic basis of these associations and the extent that these strategies may be linked through pleiotropy or correlated selection (73, 79).

For many native plants like waterhemp, adaptation to fluctuating climate must occur alongside the conversion of their habitat to agriculture. In cropping systems, high plant densities and soil compaction can reduce water availability, thereby intensifying selection from drought in largely unirrigated landscapes such as the Midwestern Corn Belt (80, 81). Our selection experiment demonstrates that agricultural populations tend to be better adapted to water limitation than natural populations, surviving longer under drought. Genomic time series analysis further resolves both a difference in drought allele frequency among habitats and stronger selection on drought alleles through time in agricultural compared to natural habitats. However, both these phenotypic and genomic dynamics could result from either increased drought pressure in agricultural systems or reduced drought selection in natural populations, which are often riparian habitats with high water availability. Given var. *rudis*’ historical association with disturbed habitats, Sauer suggested it may have been better adapted, or even “preadapted”, to agricultural habitats (40), a hypothesis supported by recent phenotypic and genomic work (20, 41). Notably, all 43 loci with protective effects under drought in this study were derived from var. *rudis* ancestry (**Fig. 4A**) and are enriched in agricultural populations beyond genome-wide expectations (**Fig. S16**). This genomic pattern mirrors our experimental results, where *rudis*-ancestry individuals and agricultural populations showed longer survival under drought (**Fig. 2**). Although we have yet to test the role of disturbance in preadaptation to agriculture, the results here, along with divergent climate histories across ancestries (82) suggest that a history of selection from drought has been particularly important in var. *rudis*’ successful invasion into agriculture.

Waterhemp’s dynamics of drought adaptation reinforces a fundamental tension: while populations can rapidly evolve under extreme selection, heterogeneity across space and time buffers these responses over larger spatiotemporal scales. Our experiments demonstrate the substantial standing genetic variation that facilitates rapid adaptive responses to drought, in part due to fluctuating selection preventing its erosion. As climate change accelerates and drought intensifies, whether directional selection will overwhelm this heterogeneity needed for climate tracking remains an open question. Ultimately, an understanding of these multi-scale dynamics across species will be critical for predicting evolutionary outcomes under accelerating environmental change.

## Acknowledgements

We thank Tyler V. Kent, Sally P. Otto, and Maryn Carlson, along with the Rieseberg, Carley, and Kreiner labs, for feedback throughout this project. We appreciate UBC Horticultural Greenhouse staff for help with experimental set-up and implementation, and undergraduate research assistants Camila Cordova and Vicky Lim for assistance with data collection. Rozenn Pineau was supported by a Dropkin Fellowship in Plant Biology at University of Chicago.

## Data Availability

Data and materials availability: All new sequence data (contemporary dataset) have been archived at the NCBI Sequence Read Archive, BioProject ID PRJNA1381560. Sequence data from the previously published herbarium dataset can be found under BioProject ID PRJNA878842. Scripts and details on analyses can be found on the github repository: https://github.com/rozenn-pineau/drought-project/tree/main/.

## Methods

### Drought selection experiment

This experiment drew from collections of 34 *Amaranthus tuberculatus* populations, sampled in design pairing nearby natural and agricultural habitats across the Midwestern United States (17 pairs total; Kreiner et al. 2021). We used seeds from 490 maternal lines from a subset of 24 populations (12 agricultural, 12 natural; **Fig. 1A**). For each maternal line, two replicates of 10 seeds were sown into separate soil discs in a growth chamber under a fluctuating temperature regime (28°C day / 18°C night) for 10 days, providing one replicate for each of two treatment groups.

Seedlings were then transplanted into 1L pots filled with Sunshine Mix #4 (Sun Gro Horticulture) and moved to the University of British Columbia agriculture greenhouse. With 490 lines x 2 treatments, the experiment included 980 pots/individuals in total. Pots were distributed in a randomized block design across two greenhouse benches. The experiment was conducted from July through August. Day length matched the natural photoperiod during this time, but full-spectrum lights were activated in the greenhouse during cloudy periods throughout the day. Late-emerging seedlings were thinned to ensure a single focal plant per pot.

Water was delivered to control and treatment pots via automated dripper lines, supplying 100 mL every 6 hours (400 mL/day) for the first three weeks of growth. During this period, two leaves per plant were collected for later DNA extraction. At three weeks, irrigation lines to all drought treatment pots were shut off completely. Stem width and plant height were measured for all plants immediately before drought onset and again 2 weeks post-treatment both for calculation of growth rate and to control for biomass-driven variation in water demand. Plant survival was monitored daily, and individuals were considered dead at the point of full wilting—defined as complete loss of turgor pressure in apical and leaf tissues (**Fig. S1)**. When this occurred, the date of death was recorded, and the individual was removed from the experiment. This design enabled us to quantify both drought avoidance and early growth rate, facilitating tests for trade-offs between these traits across natural and agricultural populations.

### Phenotypic Analysis of Drought

To analyze variation in drought avoidance in our experiment, we used individual-level mixed effects models with the structure: days to full wilt ∼ habitat + longitude + latitude + stem width + height growth rate + (1 | Bench/Tray), where nested random effects account for experimental bench and tray. Models were fit using *lmer* from the lme4 package with p-values calculated via *lmerTest*.

To further assess variation in population-level drought resistance, we calculated the proportion of surviving samples for each population for each day of the drought experiment. These population survival curves were constrained to start at 100% survival on day 1 and 0% on day 20 when all individuals had died (**Fig. 1B**). To model the dynamics of survival through time, we fitted survival curves with a logistic function using the *nls* and *SSlogis* functions in R (see **Fig. S2** for example and equation). From the estimated parameters, we calculated the lethal dose, defined as the day when 50% of the population had died (LD_50_). To quantify predictors of phenotypic variation in drought avoidance, we implemented a multiple regression model of LD_50_ against habitat, longitude and latitude: LD_50_ ∼ habitat + longitude + latitude. Model comparison using AIC indicated that the model excluding latitude provided a better fit; latitude was therefore removed from this model (**Table S1**). The partial R^2^ for each variable of interest was estimated by comparing the reduced model multiple R^2^ from the full model multiple R^2^. Additionally, we applied a non-parametric test of whether LD_50_ differed between habitats using a permutation approach. We permuted 10,000 times the habitat assignment for each population and calculated the difference between the mean LD_50_ in agricultural versus natural habitats (**Fig. S3)**. In addition, to test for the effect of habitat on drought resistance, we performed a one-sided Wilcoxon rank test with paired samples with *wilcox.test* function in R (**Fig. 1D**).

### Investigations of Trade-offs with Drought Avoidance

To test whether drought avoidance showed evidence of trade-offs with growth rate or herbicide resistance, we compared traits at the individual-level and population-level, in univariate and multivariate models (**Fig. S4**). The day of full wilt was used for individual-level analyses, and drought LD_50_ for population-level analyses. We quantified growth rate by measuring plant height before the drought treatment, along with one week and two weeks after in both control and drought-treated plants. We estimated resistance to glyphosate herbicides by quantifying depth at the EPSPS locus, as in (66). In brief, for each sample, we calculated depth within the EPSPS gene using *mosdepth* (83), which we then normalized by dividing by the mean genome-wide depth at coding sites.

We first tested whether growth rate predicted drought avoidance by comparing a maternal line’s day to full wilt in the treatment as compared to growth rate in experimental control in a univariate model and with Pearson’s correlation coefficient (**Fig. S4**). We also looked at the pairwise relationships between EPSPS copy numbers in the same fashion. We then implemented a multivariate mixed model (function *lmer* in R) of drought avoidance at the individual-level with the following structure: drought avoidance ∼ growth rate + EPSPS copy number + longitude + latitude + habitat*mean ancestry + (1|Bench/Tray). To test for population-level differences between habitats in multivariate trait space, we performed one-sided Wilcoxon rank tests (*wilcox.test* function in R) on mean drought avoidance, mean herbicide resistance, and mean growth rates, leveraging the paired sampling design (**Fig. 1D & Fig. S5**).

### DNA collection and sequencing

Genomic DNA was extracted from 300 samples using a modified CTAB protocol (84, 85). Briefly, frozen tissue was ground and subjected to a two-step extraction with CTAB-free buffer followed by 3% CTAB buffer supplemented with β-mercaptoethanol. After chloroform extraction, DNA was precipitated with isopropanol, washed with ethanol, and treated with RNaseA. The purified DNA was quantified using a Qubit fluorometer, with 280 samples yielding sufficient high-quality DNA (≥1 ng) for library preparation following the drought treatment. Illumina paired-end libraries were constructed using a protocol largely modified from (86), the TruSeq DNA Sample Preparation Guide from Illumina (Illumina, San Diego, CA) (87). Genomic DNA (1 ng) was sheared to approximately 350 bp using a Covaris ultrasonicator with the following settings: Peak Incident Power 50, Duty Factor 20%, Cycles per Burst 200, Treatment time 65 s, at 20°C. Size selection was performed using a two-step SPRI bead purification protocol with home-made SPRI beads (87), optimized to select for 350 bp fragments.

The size-selected fragments underwent end repair using NEBNext End Repair enzyme mix, A-tailing with Klenow Fragment (3’→5’ exo-), and adapter ligation with NEB Quick Ligase using P1 and P2 adapters (with unique barcodes for each sample). Libraries were enriched by PCR amplification using KAPA HiFi HotStart ReadyMix with 8 PCR cycles. The amplified libraries were purified using SPRI beads and eluted in Tris buffer (pH 8.0, 10 mM). Library quality was assessed using a Qubit fluorometer (High Sensitivity kit) and fragment size distribution was verified on a Bioanalyzer (High Sensitivity DNA chip), with successful libraries showing concentrations ≥5 ng/μl and peak sizes of approximately 470 bp (including ∼120 bp adapter sequences). Sequencing was performed at Novogene on 4 lanes of a NovaSeqX 25B. At this stage, the estimated read coverage was 11.6x for the contemporary samples, and 48.8x for the herbarium samples.

### Genome alignment and variant calling

Raw sequencing reads were processed using fastp v0.23.2 for adapter removal and poly-Q tail trimming. Quality-filtered reads were aligned to *Amaranthus tuberculatus* reference genome (haplotype 2 of accession 19_3; (88)) using BWA-MEM v0.7.17 with default parameters and appropriate read group headers, and alignments were converted to BAM format and sorted using samtools v1.15.1 with multi-threading enabled. PCR and optical duplicates were identified and marked using Picard MarkDuplicates v2.27.4. Read processing procedures were as above for historical herbarium samples, but with three modifications to account for DNA degradation. DNA damage patterns were assessed and corrected using mapDamage (89) with the --rescale option to account for typical cytosine deamination patterns in historical samples, paired-end reads were merged with fastp due to small fragment size distributions, and duplicates were called with a program DeDup which is optimized for aDNA (90). At this stage, the read coverage was on average 10.4x for the contemporary samples, and 9.7x on average for the herbarium samples.

We called genomic variants using FreeBayes v1.3.6 in parallel across 100kb genomic regions, separately for the contemporary and herbarium sample datasets. FreeBayes was run with the --use-best-n-alleles 2 and --report-monomorphic flags to call both variant and invariant sites. Variant calling was parallelized across multiple compute nodes using GNU Parallel. Raw variants were processed separately for SNPs and invariant sites using bcftools. For both site types, we applied an initial filter requiring ≤25% missing data across samples (F_MISSING <= 0.25). SNPs underwent additional quality filtering including: minimum variant quality score ≥30 (QUAL >= 30), allele balance between 0.25-0.75 for heterozygotes or ≤0.01 for homozygotes (AB >= 0.25 & AB <= 0.75 | AB <= 0.01), presence of reads on both forward and reverse strands (SAF > 0 & SAR > 0), minimum mapping quality ≥30 for both variant and reference alleles(MQM >=30 & MQMR >= 30), and paired-read support criteria. Only biallelic SNPs with allele frequencies between 0 and 1 were retained for downstream analyses.

### Genome-wide population structure and its relationship to drought avoidance

We used PLINK (version 1.9, 2024.04.1+748) (91) to perform a PCA of genotypes on a thinned set of genome-wide SNPs (**Fig. 2A**). Linkage Disequilibrium (LD) thinning was performed by calculating LD between each SNP in 100-kb windows across the genome, using a step size of 10kb, and removing SNPs with an LD (r^2^) >0.5. We detected two outlier individuals on our initial PCA that formed a very distinct group from the rest of the samples. We speculated that they might be another species of *Amaranthus* and therefore excluded them from all subsequent analyses. We used ADMIXTURE (version 1.3.0) (45) to calculate global ancestry by sample, testing K=1-6 (**Fig. 2A, Fig. S6B&C**). Multivariate linear regression models were implemented to test the effects of geography and environment on PC1, PC2: PC1/PC2 ∼ latitude + longitude + environment. We calculated the proportion of var. *rudis* ancestry by individual and population based on the genomic proportion derived from the ancestral grouping with higher frequency in the southwest at K=2. Collinearity between PC1 and ancestry proportion at K=2 (ancestryprop_K=2_) was estimated using *corr.test* in R (**Fig. S7**).

To test the effects of genome-wide ancestry (in addition to geographic, and environmental predictors) on the population-level drought LD_50_ estimates, we extended our multiple linear regression models to include ancestry as a covariate: drought LD_50_ ∼ longitude + latitude + habitat + longitude*ancestryprop_K=2_ (**Fig. 2C**), or using PC1 instead of ancestryprop_K=2_. Least square means of the effect of ancestry by longitude on drought LD_50_ was inferred from this model with *lsmips* in R (**Fig. S9**). We also performed the same models at the individual-level, testing the effect of geography, environment and ancestry/PC1 on survival: day to full wilt ∼ latitude + longitude + environment + PC1 + longitude:PC1 or ancestryprop_K=2_ (**Fig. S8**). As above, partial R^2^ for variables of interest was calculated by subtracting the multiple R^2^ value of a reduced model without ancestry as a covariate from the multiple R^2^ value for the full model.

### Local ancestry inference

To infer ancestry at each position in the genome, we used AncestryHMM (92). This approach requires (1) the allele counts for each of the parental ancestry panels (var. *rudis* and var. *tuberculatus*), (2) the recombination distance between each locus, and (3) the genotype call for each focal sample at each ancestry informative site.

To generate allele accounts for the ancestry panels (1), we used an independent and previously generated dataset (41) selecting pure var. *rudis* and var. *tuberculatus* populations for the two ancestry panels. To determine which individuals were of pure ancestry, we mapped reads to the newest reference genome version (88) and called and filtered SNPs as above before running ADMIXTURE (version 1.3.0) assuming k = 2. We identified 44 pure *var. rudis* and 21 pure *var. tuberculatus* samples using a threshold of 0.00001 alternate ancestry. We filtered the variant file (50,811,811 SNPs) for this set of samples and applied a minimum allele frequency cutoff of 0.05. To identify ancestry-informative sites, we calculated F_ST_ between ancestry types using VCFtools (version 0.1.16) on a site-by-site basis, and retained variants as ancestry informative if their F_ST_ fell above the 75th percentile (2,089,620 variants). We intersected these sites with SNPs from the drought selection experiment, resulting in 788,177 markers for inference of fine scale ancestry (∼ 1/kb). Finally, we extracted population-level allele counts for each ancestry grouping for each site using a custom bash script.

To generate the linkage map for (2), we used a set of scripts https://github.com/QuentinRougemont/LDhat_workflow using the interval algorithm. We used a precomputed look-up table based on theta = 0.01, n = 100. We then estimated *r* between SNPs from the ldhat output based on rho = 4N_e_*r* with N_e_ = 540,804 as estimated in (66). Then, we fit a monotonic spline on the cumulative distribution of these recombination rates, to calculate the physical distance along the chromosome and between each ancestry informative SNP in the drought variant call set presented in (1). Finally for step (3), we extracted the read count for each ancestry informatic site for each focal sample in the drought dataset from the filtered variant file using a custom bash script (788,177 variants and 280 individuals).

We implemented ancestryHMM with overall ancestry proportions of 33% of var. *tuberculatus* and 67% *var. rudis*, based on the mean of ADMIXTURE estimates across our accessions. We modelled two pulse events that introgressed *var. rudis* into *var. tuberculatus* with a prior of 10,000 and 100 generations ago. The resulting model predicted an initial pulse of var. *rudis* into var. *tuberculatus* of 42.54% 10,000 generations ago, followed by a more recent pulse of 24.56% 7.45 generations ago. We assigned ancestry based on probability outputs from the model with a cutoff of 0.9 (NA below that threshold). Probabilities were given for how many *var. rudis* allele was derived from pulse one or two. Since we were only interested in the ancestry genotype (rather than pulse-specific origin), we summed across pulses to obtain the count of *var. rudis* alleles (0, 1, or 2) for each individual at each locus. Collinearity between the individual mean fine scale ancestry composition and ADMIXTURE estimates was then compared (**Fig. S11)**.

### Admixture mapping

To detect significant associations between genome-wide loci and drought avoidance, we first conducted a Genome Wide Association Study (GWAS) with the day to full wilt/survival under drought as the dependent variable. Since drought avoidance is highly correlated with ancestry, we expected the risk of false positives with a SNP-based GWAS while controlling for structure to be high (**Fig. S10**). To address this, we conducted an admixture mapping approach leveraging fine-scale ancestry estimates, after filtering out loci with > 20% missing data. We used GEMMA (v0.98.5), to associate day to full wilt with ancestry calls across the genome, controlling for population structure with the mixed-effect model implementation (−lmm) using a relatedness matrix based on ancestry calls. We applied a genomic inflation factor (λ = 2.35) to correct for an overrepresentation of significant variants with low - log10(p-values) (**Fig. S12**). In all cases, qqplots used to investigate the distribution of significance were produced with the r package qqman (**Fig. S12)**. FDR correction was then applied to these *p*-values, with a threshold of q = 0.05 as indicated in **Fig. 3B**.

To identify independent drought-associated variants, we performed LD-clumping on SNPs that passed the q = 0.05 threshold. To do so, we used plink --clump, dropping SNPs with an r^2^ > 0.5 with a focal index SNP within a 100 kb window. We used bedtools intersect to identify the genes that drought-associated SNPs mapped to. Using both the de novo annotation file and *Arabidopsis* orthologues identified with orthofinder2 (93), we identified a set of genes for GO enrichment analysis using PANTHER 19.0 (94) **(Table S6**). To link genes to function, we searched the literature for evidence of drought implication of the most important SNP hits in our dataset (**Table S5**). We also used snpEff (95) with a custom *Amaranthus* database, to identify the type and effect of mutation on the protein sequence.

To estimate LD decay, we first filtered the genotype dataset to retain SNPs with a minor allele frequency ≥ 0.05 and present in at least 80% of individuals. To reduce computational burden while preserving LD structure, SNPs were further thinned by removing one SNP every 2 kb. LD decay was then calculated using PopLDdecay (96). An exponential decay model was fitted to the mean *r^2^* values as a function of physical distance, *r^2^ = a * exp^−bd^ + c*, where *b* represents the LD decay rate, which also allowed us to solve for the half decay distance (**Fig. S13)**.

### Relating Adaptation to Agriculture to Drought Avoidance

To identify loci with consistent divergence between habitats among replicated population pairs, we implemented a Cochran-Mantel-Haenszel (CMH) test with plink v1.9 (91) on genome-wide SNPs from the contemporary drought sequenced samples (as in (20)). We selected a relatively arbitrary FDR threshold of 0.01, which yielded 94,008 SNPs surpassing this multiple test correction cutoff, representing SNPs enriched for a role in adaptation to agriculture (**Fig. S15**). To test whether there was a significant enrichment of drought-adapted loci in this set of putatively agriculturally-adapted loci, we compared the observed number of CMH SNPs falling within the set of bounds of the 43 drought-adapted clumps compared to a null based on the distribution of haplotype sizes. To generate the null, we first calculated the ancestry block length spanning each of the 43 lead SNPs and for each sample. Based on the mean ancestry tract length across samples at each lead SNP, we calculated the number of CMH SNPs falling within those ancestry block boundaries. To generate a null distribution, we then used a permutation approach where: we randomly sampled 43 loci with similar ancestry block mean lengths, and calculated the number of CMH SNPs falling within those randomly sub-sampled ancestry block boundaries. This process was repeated 1,000 times, with two stringency levels when subsampling similar-sizes loci, 1kb and 10kb (**Fig. S14**).

To test whether drought alleles were particularly enriched in agricultural compared to natural habitats, we first estimated individual level drought-allele frequency by averaging genotypes across the 43 previously identified drought-associated loci. We fitted a linear model predicting the collective frequency of these drought-associated ancestry tracts as a function of environment, latitude, longitude, and their interactions, while controlling for genome-wide ancestry proportions (as inferred at K = 2 from ADMIXTURE) as a covariate. Model significance was assessed using Type III ANOVA. To visualize significant interaction effects, we generated least-squares means predictions across the range of observed latitudes (38.5–41.5°N) and longitudes (83–95°W) (**Fig. S16**), calculating the difference in predicted drought allele frequency between agricultural and natural populations at these geographic extremes.

### Within-generation survival-based selection coefficients (s_β_), allele frequency change, and Cox hazard analyses

To determine the strength of selection in our drought experiment, we fit a logistic model to the daily change in individual haplotype frequency during the drought survival experiment. To do so, we first calculated daily haplotype frequencies using genotype assignments from the ancestry mapping analysis, combined with the survival data from the drought experiment. Genotype value was rescaled to allele frequency, encoded as 0 (var. *tuberculatus* homozygote), 0.5 (heterozygote) or 1 (var. *rudis* homozygote) (**Fig. 4A**). Missing genotypes (./.) were excluded from frequency calculations. To fit a logistic model to these allele frequency trajectories through time, we further scaled individual genotypes by whether or not an individual was observed on each day. Critically, individuals were coded as missing (NA) for all days following their day they died to avoid artificially deflating selection coefficients by including dead individuals in frequency calculations. For each locus separately, we regressed this vector of individual genotypes remaining in the experiment on when they were observed using the *glm* function in R with the *“family=binomial”* option (formula: scaled genotype ∼ day). The slope of this model represents the change in logit-transformed allele frequency per day of drought treatment, reflecting differential survival favoring drought-adapted alleles. When additivity is assumed, 2 x slope = s_β_, reflecting the fitness advantage of the homozygous alternate genotype. Unlike traditional selection coefficients estimated from frequency changes across generations, this within-generation coefficient (s_β_) directly measures selection as it occurs during the drought event (**Fig. 4B**). To estimate how these coefficients changed throughout the drought, we applied this logistic regression to progressively truncated time series, stopping at each day of the experiment.

To test whether the increase in drought-adapted loci frequency over the course of the experiment was enriched relative to the genome-wide background, we implemented the following permutation procedure. We first selected a set of 43 loci that had the same starting allele frequency distribution (binned precision of 0.003) from the full set of ancestry-informative loci (>700,000 sites) as our set of 43-drought adapted loci. We calculated the mean change in allele frequency between the first and last day of the selection experiment for each set. We repeated this procedure 10,000 times to generate the null expectation of mean allele frequency change in our drought experiment (**Fig. 4C**). Statistical significance was assessed by calculating the proportion of permuted means that exceeded the observed mean frequency change.

We were interested in evaluating the collective protective effects of var. *rudis* ancestry tracts on drought tolerance, and the extent to which genome-wide ancestry would still be informative. To do so, we first constructed polygenic scores (PGS). We did so while accounting for linkage disequilibrium and avoiding effect size inflation from winner’s curse by applying a lasso regression approach (implemented in R package *glmnet*), which 1) fits all 43 clumped candidate loci simultaneously, 2) automatically selects independent signals through L1 regularization, and 3) provides shrinkage-corrected effect sizes. We performed k-fold cross-validation to identify the optimal lambda penalty parameter that minimized test mean squared error. The resulting lasso model selected 12 independent loci from the 43 genome-wide significant variants and assigned regularized effect size coefficients (β) ranging from 0.04 to 0.51. For each individual, we calculated PGS as the sum of products of genotype dosage (0, 0.5, 1) and lasso-derived β at each of the 12 selected loci. Missing genotypes were imputed using the mean genotype frequency at each locus. The resulting PGS values ranged from 0.168 to 4.33 (SD = 1.14). We then implemented Cox proportional hazards regression model of PGS using the function *coxph* in R with and without genome-wide ancestry as a covariate. Protective effects against drought per unit predictor (*k*) was calculated as 1-HR*^k^*. To obtain the Kaplan-Meier curves, we first stratified samples by their PGS and genome-wide ancestry (high/low relative to median). We then used *survfit* in R to calculate the survival curves based on the fitted Cox model.

### Spatiotemporal Genomic Analyses of Climate Adaptation

To analyze the spatiotemporal dynamics of drought adaptation, we subsampled 87 specimens from a collection of previously sequenced herbarium specimens (20), that were collected between 1880 and 2011, to match with our record of climate data. We first identified overlap between previously identified ancestry informative loci (974,134 SNPs) and the variants in the herbarium dataset using bedtools intersect. Local ancestry in herbarium samples was then inferred using ancestry_hmm with the same approach as described above. To identify the drought-adapted loci in the herbarium dataset, we compared the contemporary drought-associated loci to the herbarium loci (680/893 loci in common). We further clumped this dataset using plink (100kb-windows, r^2^ = 0.5) and obtained a set of 33 focal drought-adapted SNPs. To correlate climate to haplotype frequency change through time in the herbarium samples, we pulled data from the NOAA yearly climate database (https://www.ncei.noaa.gov/pub/data/ghcn/daily/by_year/, downloaded on 04/25/25) from across sample collection dates. We used a custom R script to identify the nearest NOAA weather station to each of our herbarium samples in the year of its collection. Since drought is associated with low precipitation and high temperatures and because *A. tuberculatus* is a summer annual, we extracted the maximum temperature (in Celsius) and total precipitation (in mm) across June, July, and August (representing climate conditions in its growing season). We removed samples where the closest weather station was more than 50 km from the samples locality resulting in 95 samples ranging from 1880 to 2011 (88 for precipitation and 59 for maximum temperature, **Fig. S17**). As in (20), habitat was assigned to herbarium samples based on annotations of habitat descriptions on the specimens.

To analyze the dynamics of drought-adapted haplotype frequency change over short timescales, we implemented an analysis that correlated change in climate to changes in genotype frequency among pairs of samples. This approach was designed to minimize the influence of spatial variation while maximizing the detection of temporal evolutionary responses to climate change. We first identified all possible sample pairs from our dataset, and ranked sample pairs by geographic distance, with the closest pairs prioritized for selection. For each drought-associated SNP, we used a greedy pairing algorithm that prioritized geographically proximate samples while ensuring non-overlapping sample usage within each SNP analysis. The pair selection process involved several filtering criteria: pairs must span different collection years (year difference > 0), geographic distance must be ≤100 km between samples (calculated using Vincent ellipsoid distance), both samples in a pair must have non-missing genotype data for the focal SNP, and only SNPs with ≥3 valid pairs were included in analyses.

For each drought-associated SNP, we calculated the Spearman’s rank correlation coefficient between temporal changes in drought allele frequency and climate between paired samples, separately for total growing season precipitation and growing season maximum temperature. Statistical significance was assessed through a permutation procedure that maintained the spatial-temporal structure of the genetic data while randomly reshuffling climate values among samples once per permutation. The same randomized climate dataset was used across all SNP analyses within each permutation. We calculated multiple test statistics including the number of significant positive correlations, the median correlation across SNPs, and the Wilcoxon signed-rank test. Permutated *p*-values were determined as the proportion of permuted datasets yielding test statistics equal to or more extreme than those observed in the observed data.

To quantify the collective trajectory of drought alleles on average over the last century depending on climate, we implemented linear and logistic multiple regression analysis. As climate has changed non-linearly through time, we were particularly interested in testing for interactions between year and climate variables (temperature and precipitation) while controlling for other covariates. We first tested a model of drought haplotype frequencies as a function of temperature, year and geographic coordinates (genotype frequency ∼ temperature max*year + precipitation*year + locus + latitude + longitude) (**Fig. S18**). We then tested the effect of adding collection habitat to the model (natural/agricultural/disturbed) (genotype frequency ∼ temperature max*year + precipitation*year + habitat*year + locus + latitude + longitude), and assessed the correlation structure between all predictors (**Fig. S19)**. Significance was assessed in all models with a type III ANOVA (*Anova* function in R). Predicted haplotype frequencies were calculated using the *lsmip* function, while the linear rates of change (β) and logit transformed rates of change (selection coefficient) for these trajectories were extracted with the function *emtrend*. We multiplied these rates by 2 to scale them as effects on homozygous genotypes (AA vs. aa) rather than single-allele effects, yielding β_AA_ and s_AA_.

## Notes

### Competing Interest Statement

The authors have declared no competing interest.

### Summary of Updates

Added permutation analyses and updated multivariate models. Added Supplementary Figures and Tables. Modified the text throughout to improve the use of terminology.

