## Supplementary Figures and Tables for "Genomic ancestry predicts rapid responses to drought across spatiotemporal scales"

**Supplementary Materials for Pineau et al., 2025**

_Supplementary Figures_

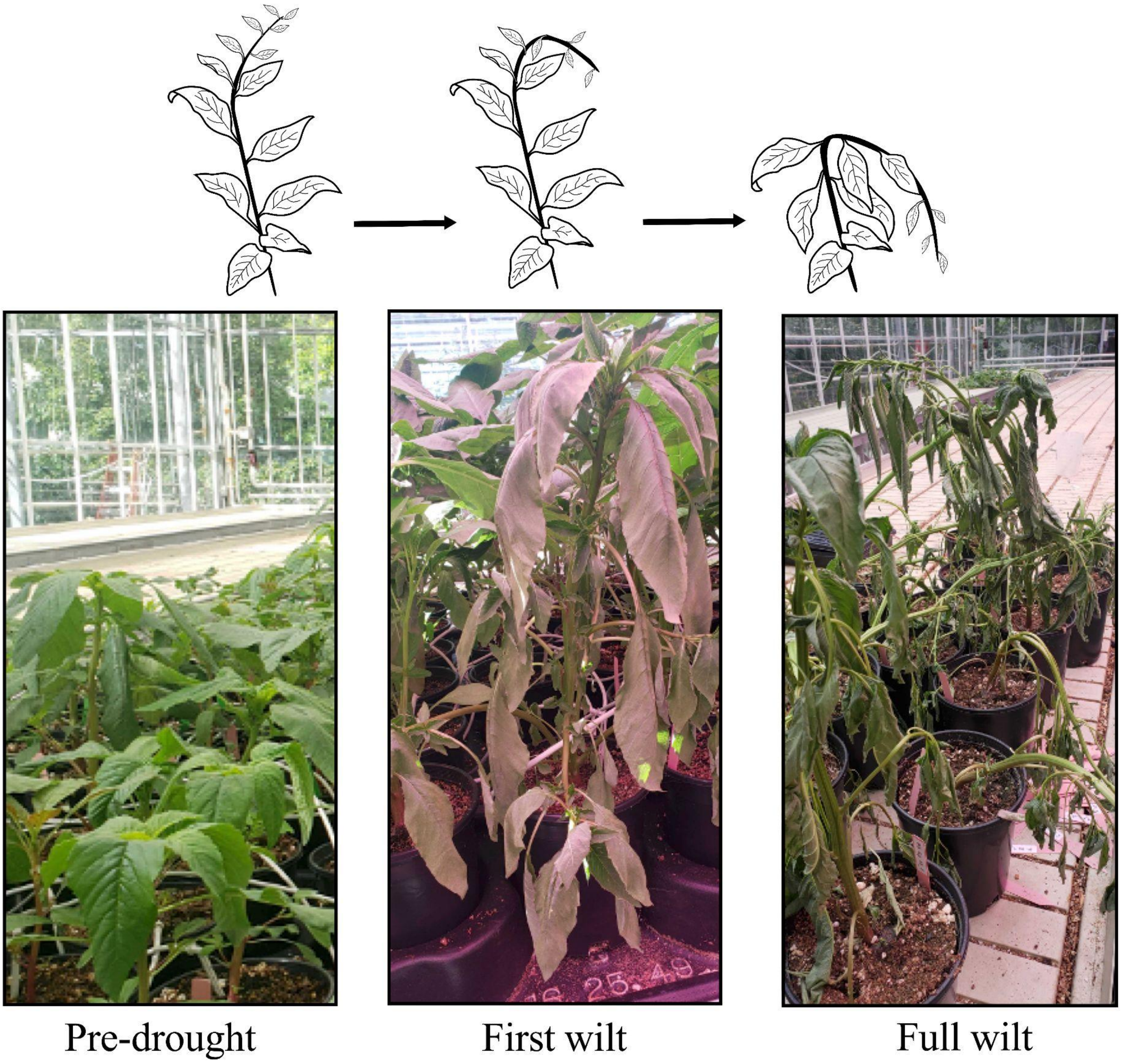

**Fig. S1.** Plant survival under drought conditions was monitored every day during 20 days (until all drought-treated plants had died). The day at which the first signs of substantial leaf wilt were observed was labeled as “first wilt”. The day to “full wilt” corresponds to complete loss of turgor pressure throughout the plant.

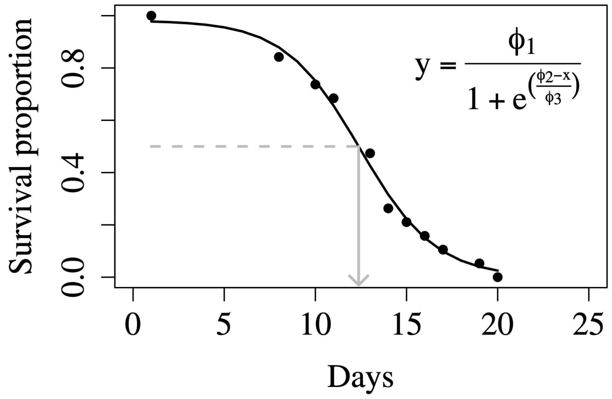

**Fig. S2.** A logistic model was fitted to the survival curves using *nls* and *SSlogit* in R. The parameters estimated from the fit were then used to calculate $LD_{50}$, the day at which half of the population has died.

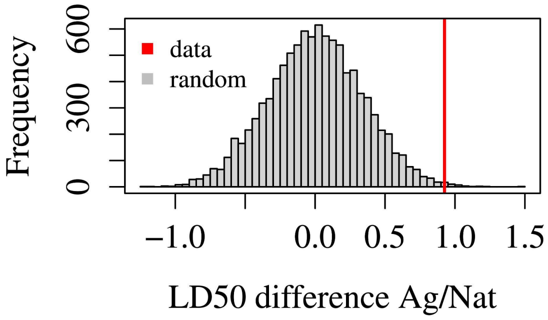

**Fig. S3.** Populations in agricultural habitats are significantly more resilient to drought as compared to those from natural habitats. We permuted 10,000 times the population assignment for their habitat, and calculated the difference between the mean $LD_{50}$ value in agricultural versus natural habitats (gray distribution). The mean value for the data is shown in red (*p*-value = 0.0036).

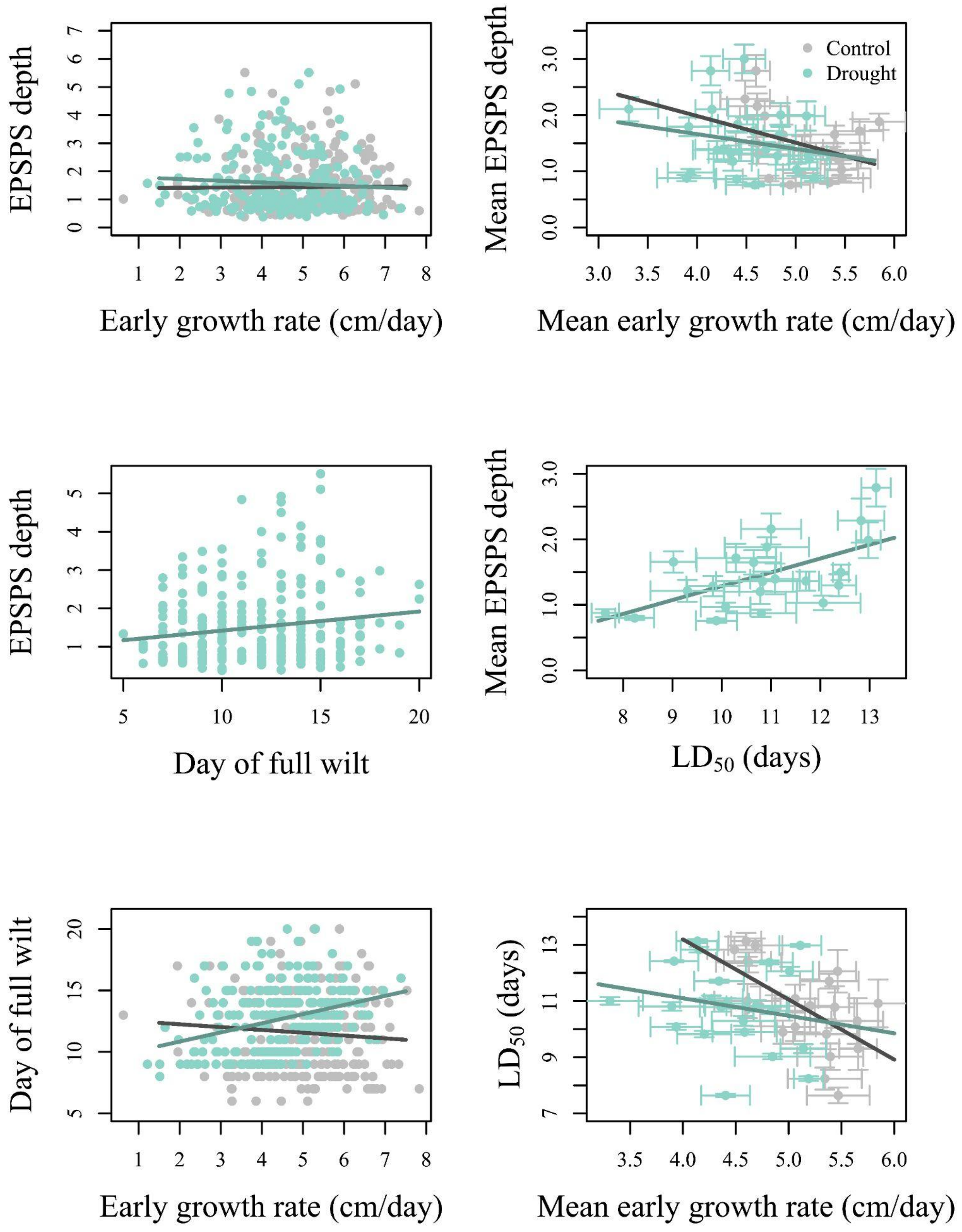

**Fig. S4.** Individual-level trade-offs (left) and population-level trade-offs (right) between growth, resistance to drought (day to full wilt and $LD_{50}$) and resistance to herbicide (EPSPS copy number). The growth rate was measured in both control (gray) and drought treated (teal) individuals. To look at the effect of drought on growth versus survival, we match the survival values (day to full wilt and $LD_{50}$) of the treated plants to their half sibling counterparts.

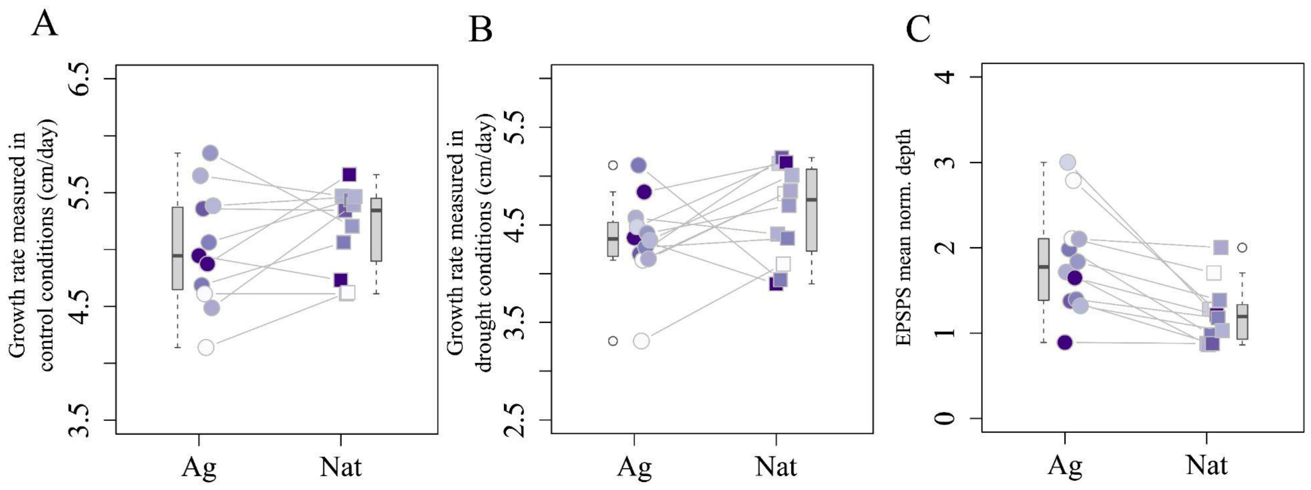

Fig. S5. Mean growth rate in control (A) and drought (B) conditions per population, and EPSPS mean normalized copy number per population (C), compared between environments (Ag/NAt). Wilcoxon paired test for growth rate in drought conditions, *p*-value = 0.06; Wilcoxon paired test for growth rate in control conditions, *p*-value = 0.89; Wilcoxon paired test for EPSPS mean normalized depth, *p*-value = 0.00024.

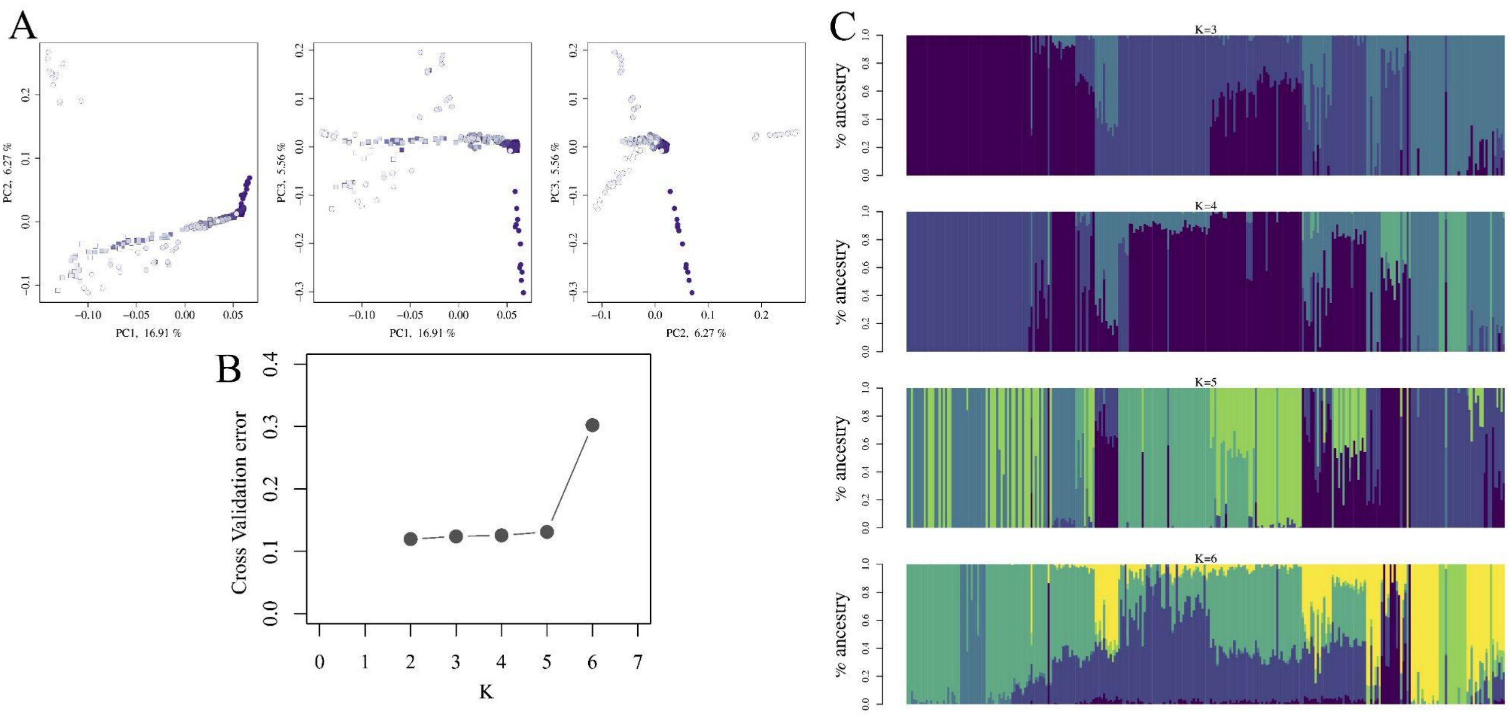

**Fig. S6. (A)** Pairwise comparisons of PC1, PC2 and PC3 axes. **(B)** Cross validation error results and **(C)** admixture bar plots from ADMIXTURE analysis for different values of K.

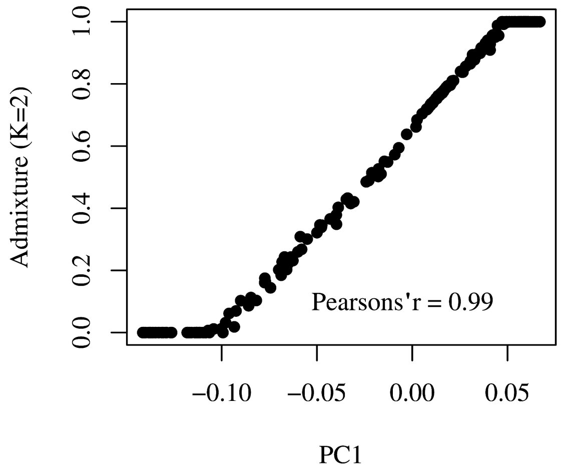

**Fig. S7.** PC1 values are highly correlated with admixture for K=2 (Pearson's r = 0.992, 95% CI = [0.990, 0.994], *p*-value < 2.2 $x10^{-16}$).

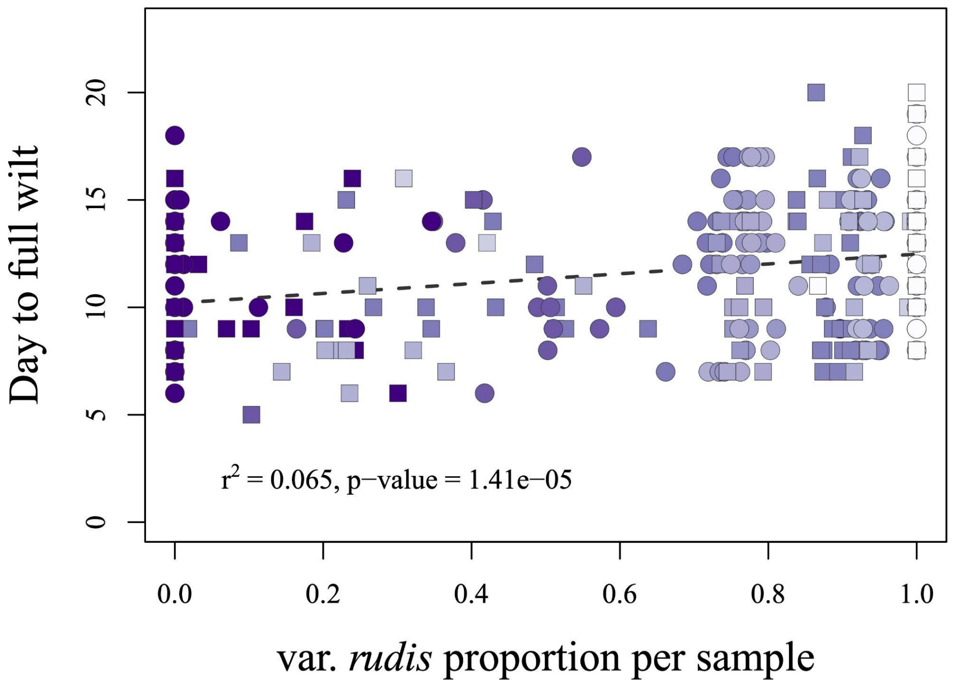

**Fig. S8.** Ancestry and longitude explain drought tolerance at the individual-level (multivariate model: ancestry: $F_{1,276}$= 36.99, *p*-value = 0.05; longitude x ancestry: $F_{1,276}$ = 43.07, *p*-value = 0.03, ancestry partial $R^2$ = 0.033, univariate model $r^2$ = 0.065). Samples are colored by longitude

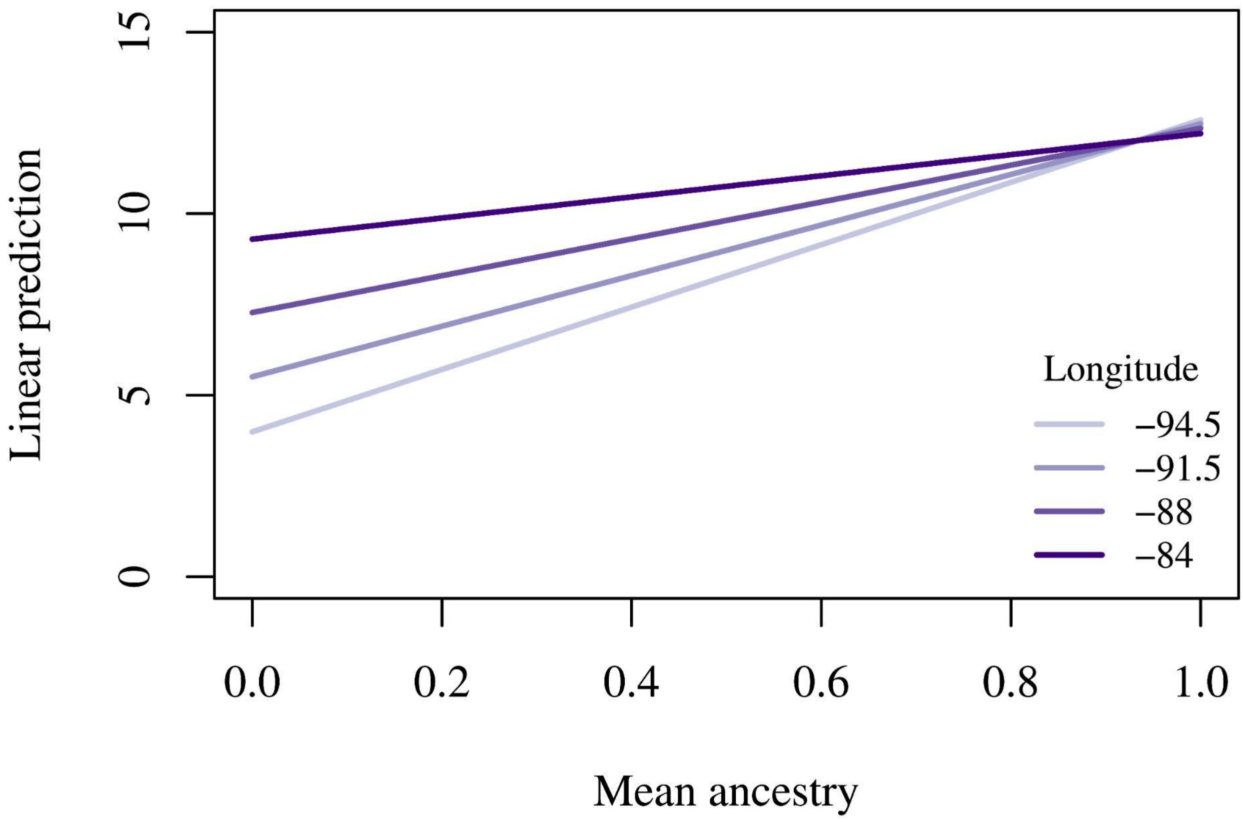

**Fig. S9.** Linear predictions of drought resistance ($LD_{50}$, in days) of the effect of admixture and longitude from a multivariate model. The combined effects of admixture and longitude were averaged over the habitats (natural and agricultural). Predictions in drought response depend on the interaction between geography and ancestry, with pure var. *rudis* ancestry being always advantageous independent of the longitude.

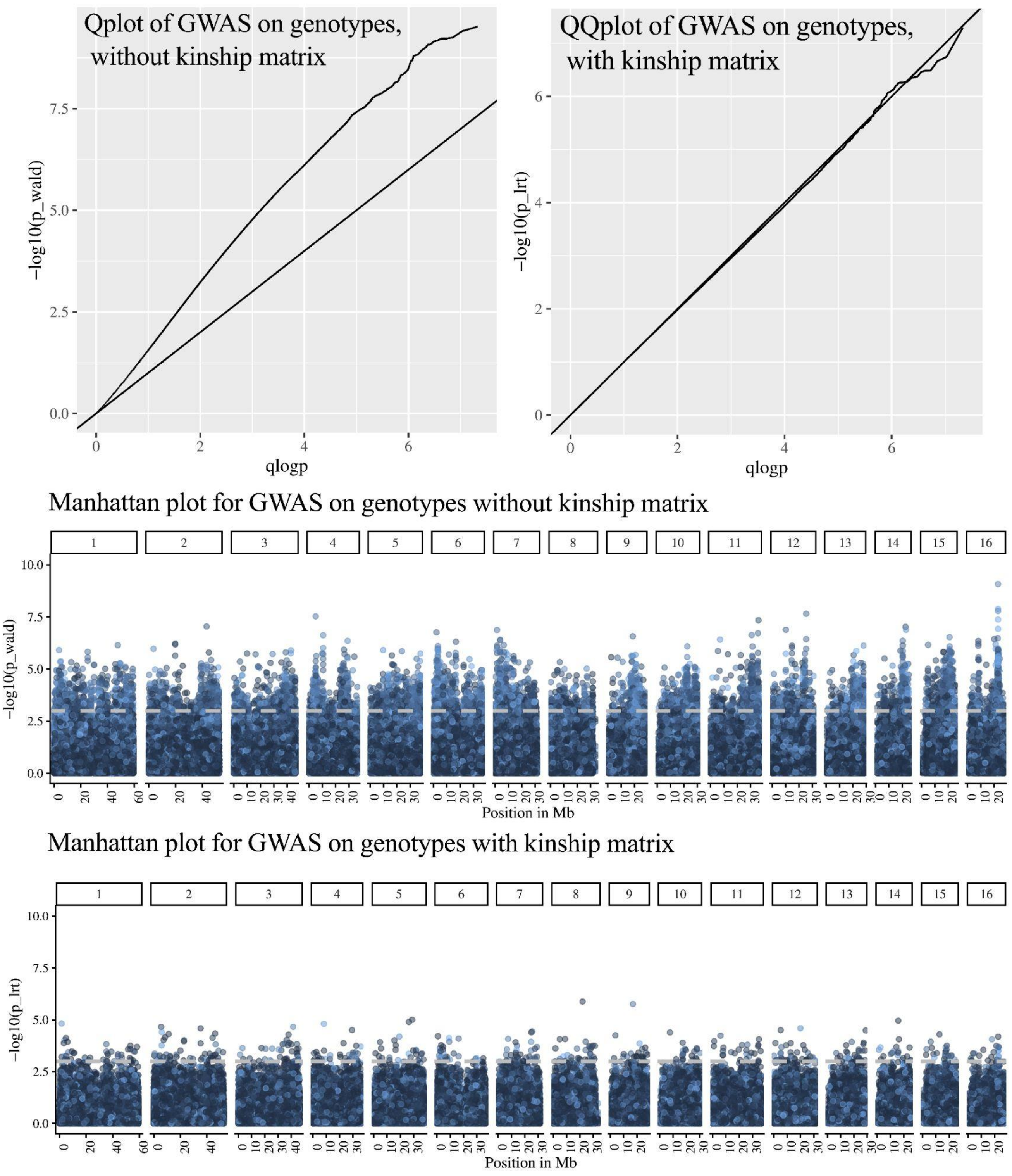

**Fig. S10.** QQplots and Manhattan plots for the GWAS on genotype values with the day to full wilt/survival under drought as the dependent variable, before and after correction for population structure with a kinship matrix correcting. Grey dotted lines represent the FDR value of 0.05.

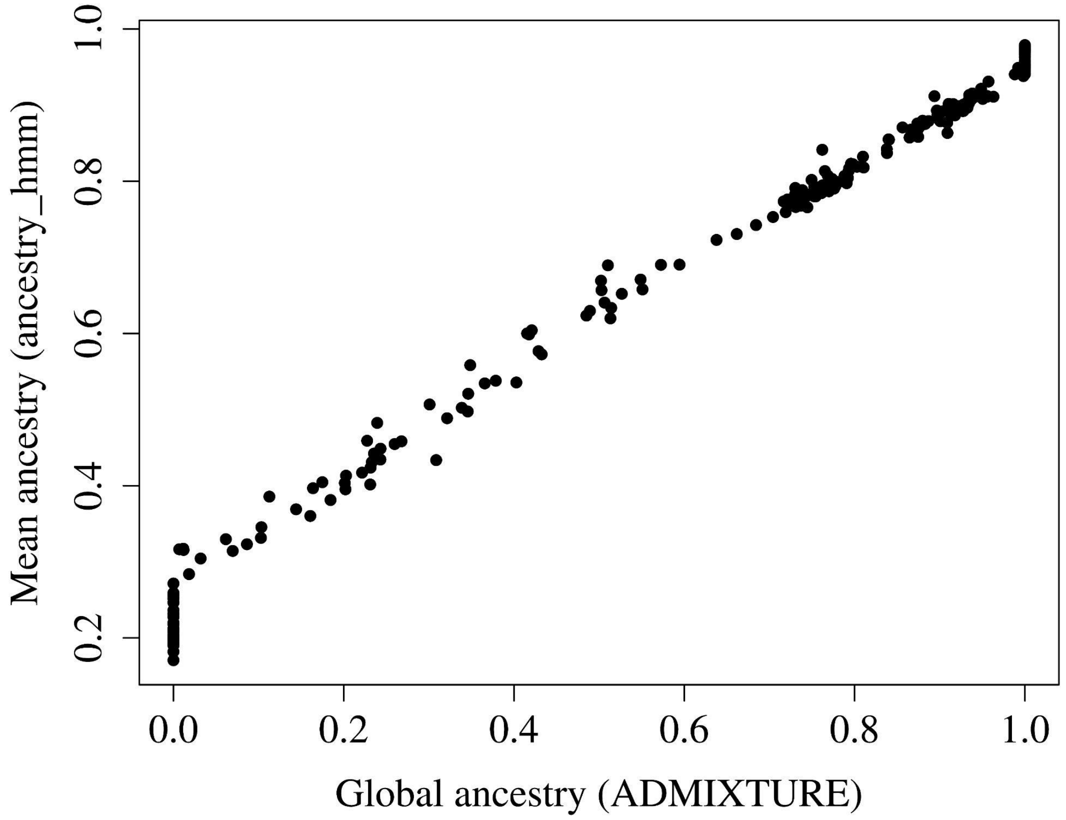

**Fig. S11.** Mean ancestry as calculated from the results of admixture mapping correlate to the global ancestry values resulting from ADMIXTURE analysis ($r^2 = 0.99$, *p*-value < 2.2 x$10^{-16}$).

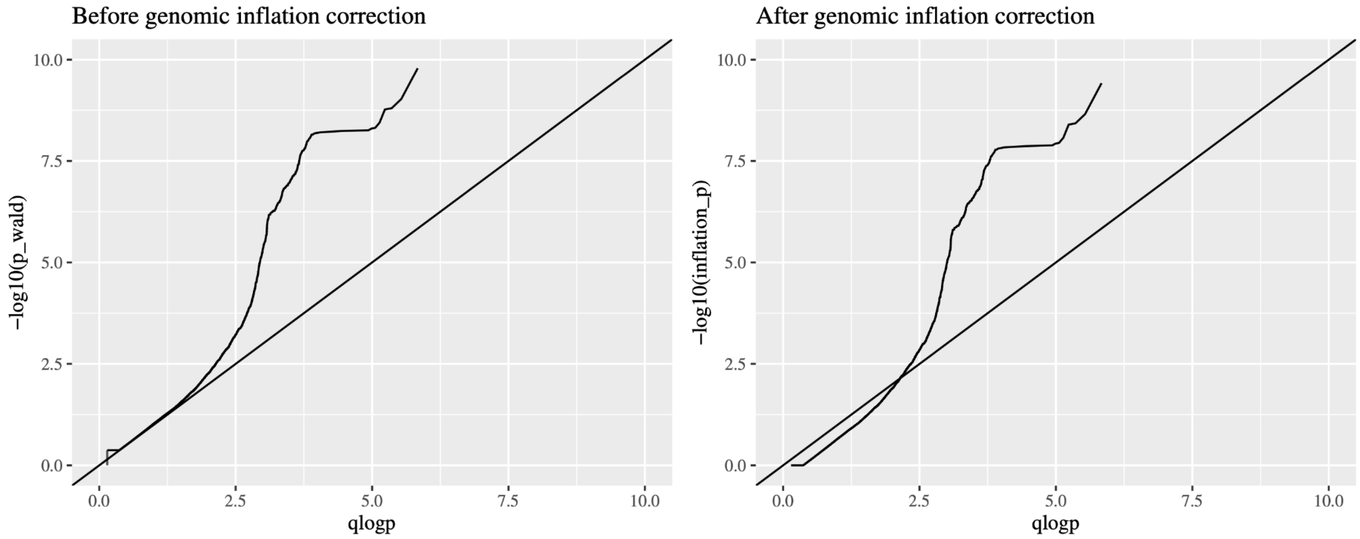

**Fig. S12**. QQplot for admixture mapping before and after genomic inflation factor correction.

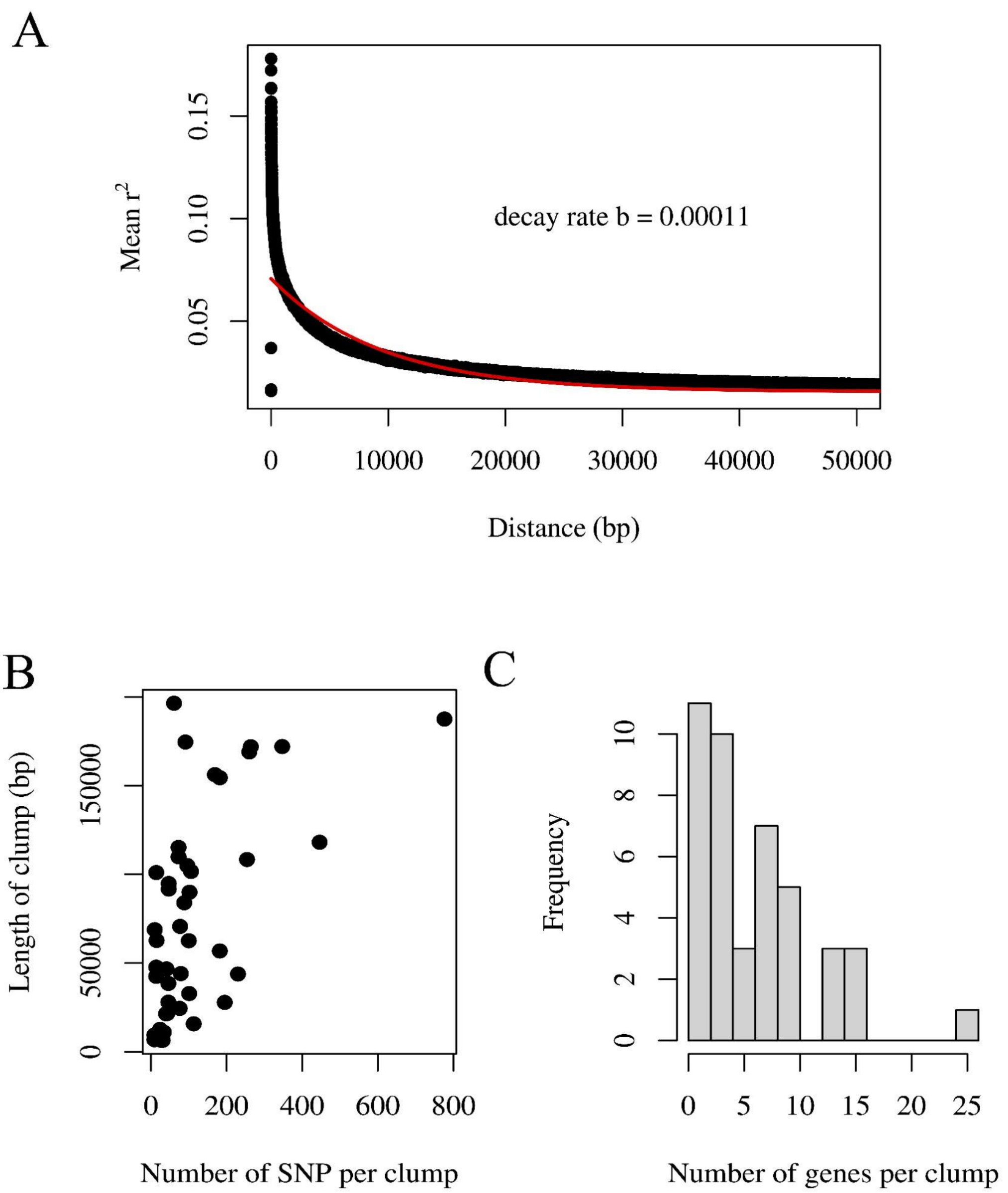

**Fig. S13. (A)** We calculated the LD decay rate on the filtered genotype file using PopLDdecay (Zhang et al. 2019). **(B)** The size of the 43 clumps with an associated lead locus ranged from 6.3 kb to 196.4 kb, with a mean of 77 kb. The number of SNPs per clump, which varied from 9 to 776 with a mean of 117.5 SNPs per clump. **(C)** The number of genes in each clump varied from 1 to 25 with a mean of 6.4.

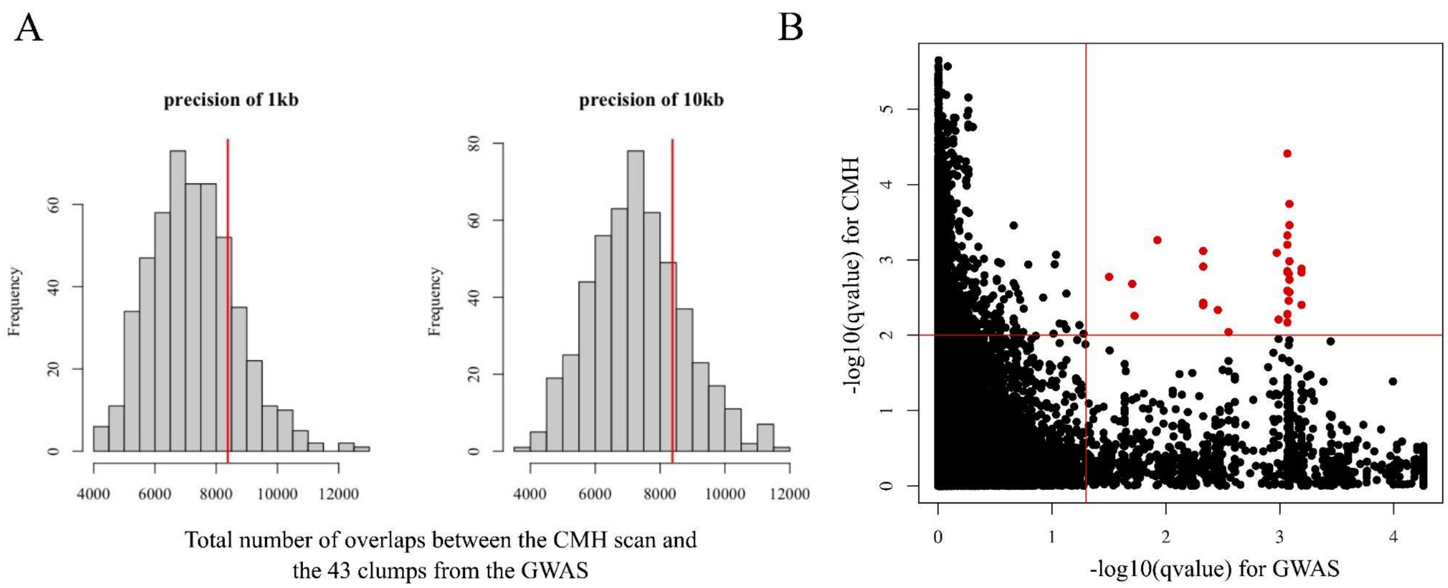

**Fig. S14. (A)** The total number of loci from the CMH scan that fall within the ancestry tracts of the 43 drought-adapted lead loci is not different than what we would expect by chance (8,381 loci, $p = 0.16$ for a precision of 1kb and $p = 0.18$ for a precision of 10kb, FDR threshold < 0.01). **(B)** Common loci (black dots) and significant loci (30 loci, red dots) from the intersection between the CMH scan and the admixture mapping on the drought dataset. Lines denote the 0.05 (admixture mapping hits) and 0.01 (CMH hits) FDR threshold we used for detecting significant variants. Genes like *HAT14, RDK5*, *TBR* (all in *Arabidopsis thaliana*) and *TDC2* (*Camptotheca acuminata*) figure among the genes in common with both admixture mapping and CHM scan significant hits (see **Table S5**).

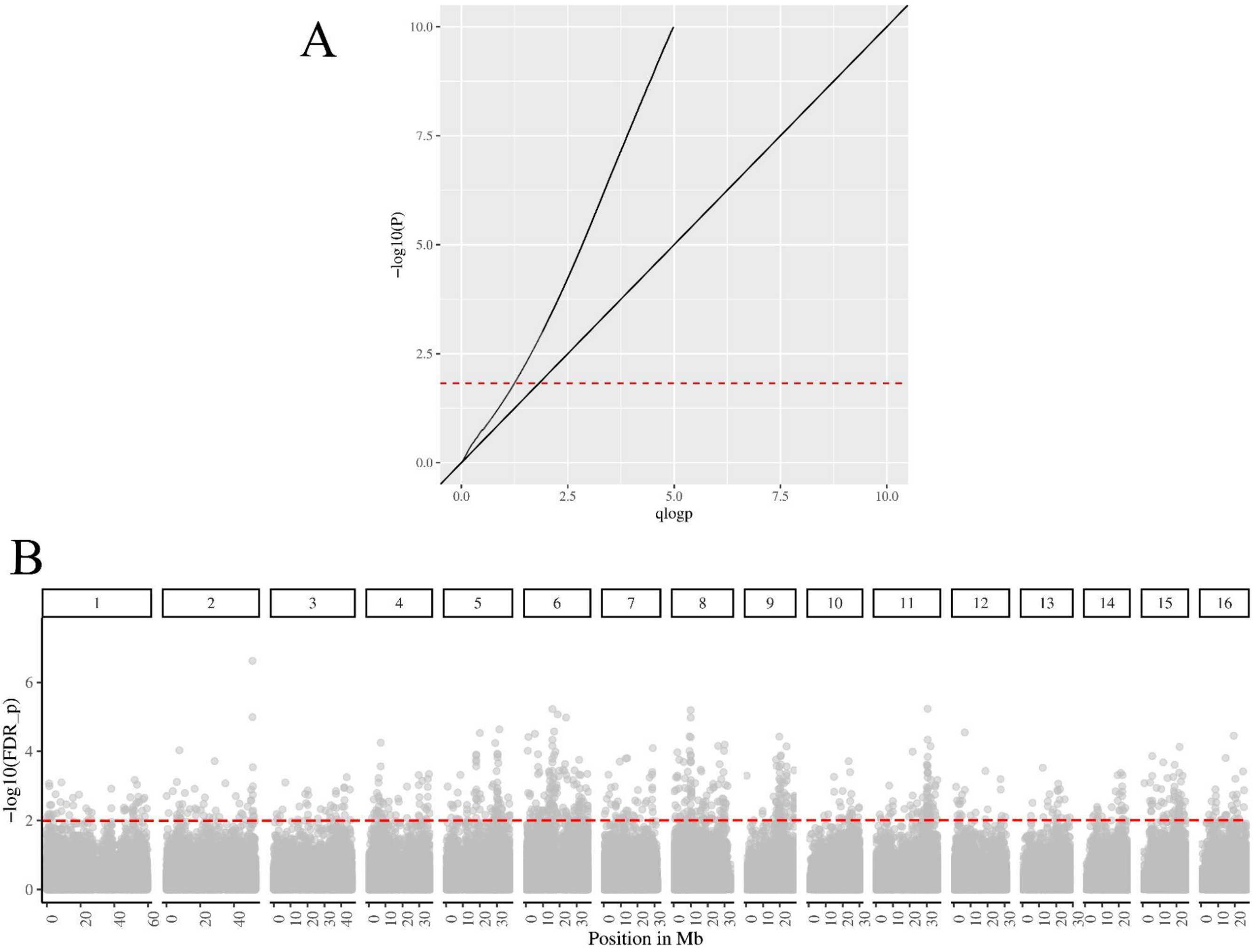

**Fig. S15**. **(A)** QQplot and **(B)** Manhattan plot for the CMH scan. Manhattan plot data points were subsampled to 200,000 SNPs for visualization purposes.

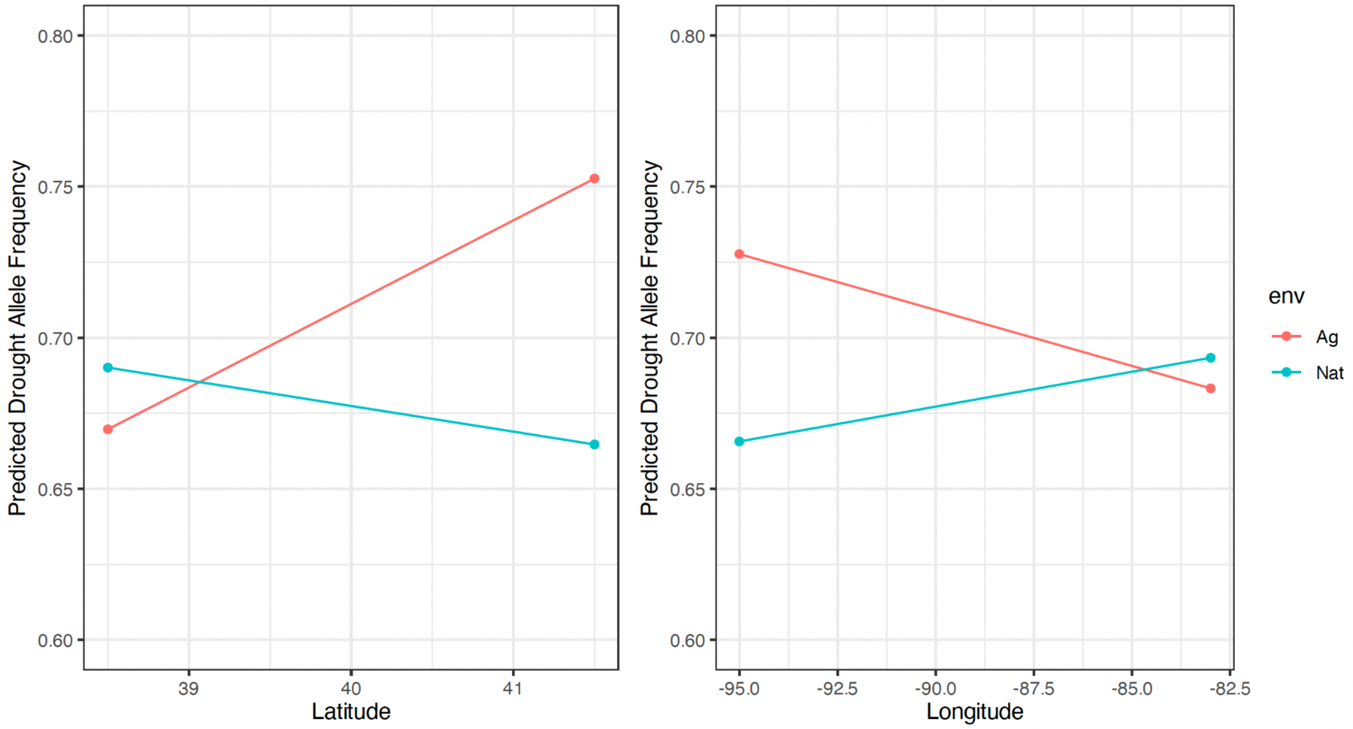

**Fig. S16.** The predicted effect of habitat on drought allele frequencies in contemporary sequenced samples from the drought experiment varies significantly with longitude and latitude, based on a multivariate linear model that controls for genome-wide ancestry.

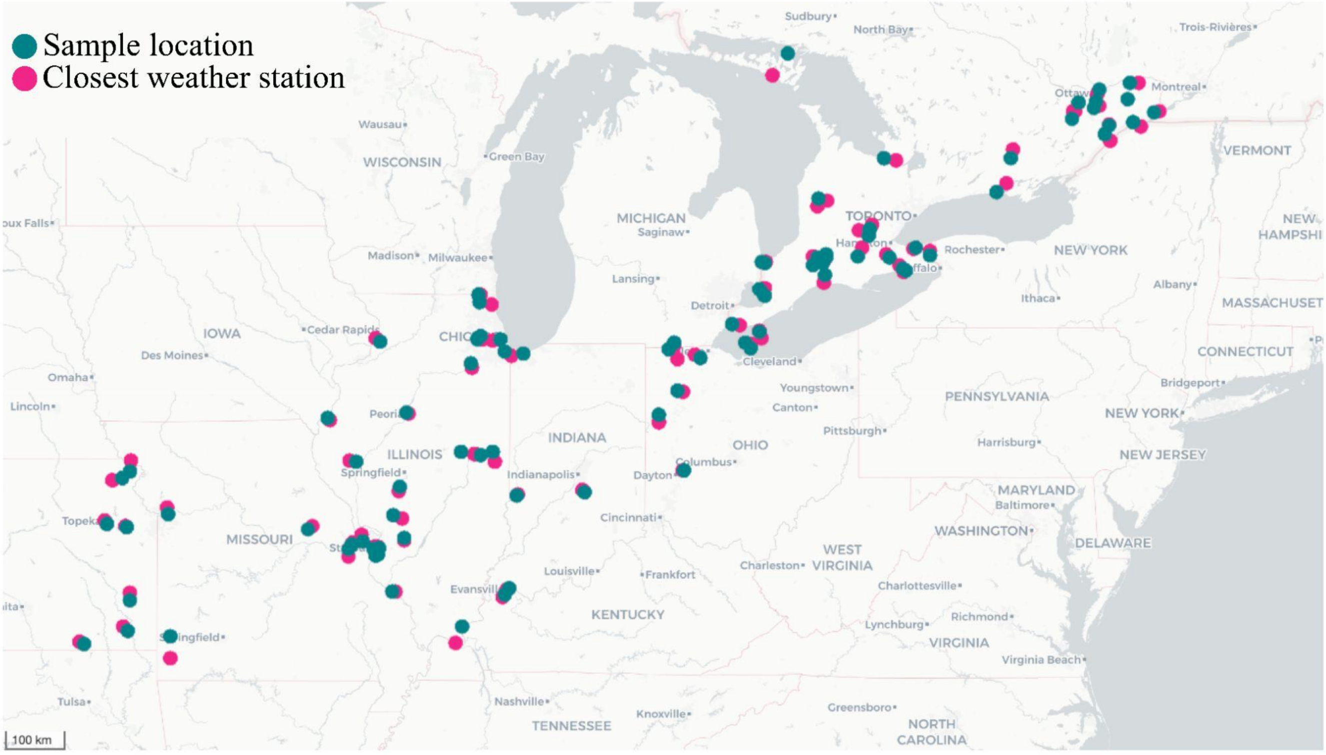

**Fig. S17.** Map of the herbarium samples (teal dots) and their closest weather station (pink dots, with a higher distance limit of 50 km).

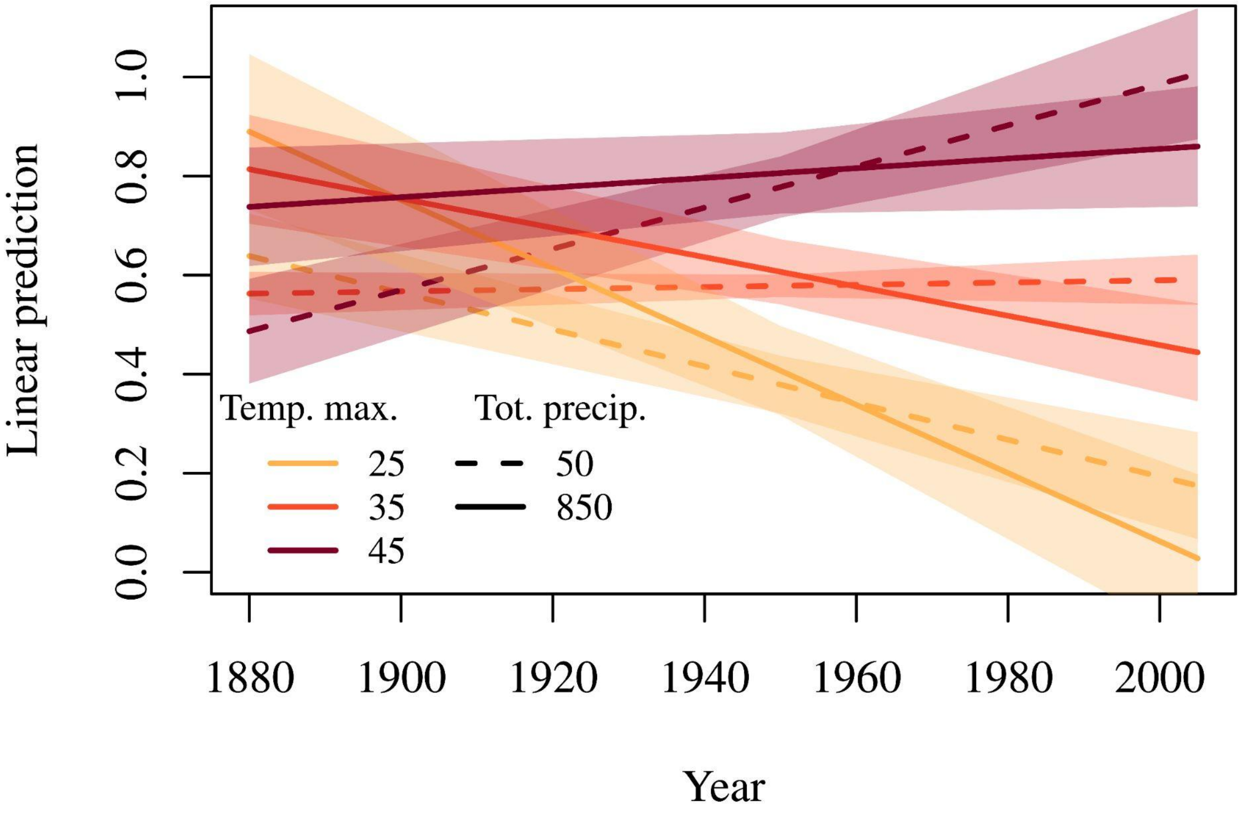

**Fig. S18**. When including only geographic and temporal predictors, the collective trajectory of drought loci (predicted allele frequency through time) is temperature (°C) dependent—with evidence of positive selection in hot years and negative selection in average or cool years ($F_{1,1088}$ = 8.70, *p*-value = 0.003 for the effect of the interaction between time and temperature, $F_{1,1088}$ = 8.41, *p*-value = 0.004 for the effect of temperature). The effect of precipitation (mm) on drought allele frequency trajectories is marginal ($F_{1,1088}$ = 3.03, *p*-value = 0.08) as is the interaction of precipitation with time ($F_{1,1088}$ = 2.98, *p*-value = 0.08). Predicted values inferred from a multivariate regression, also controlling for geographic variables along with sample year and climate. Model formulation : haplotype ~ tmax*year + prcp*year + locus + latitude + longitude.

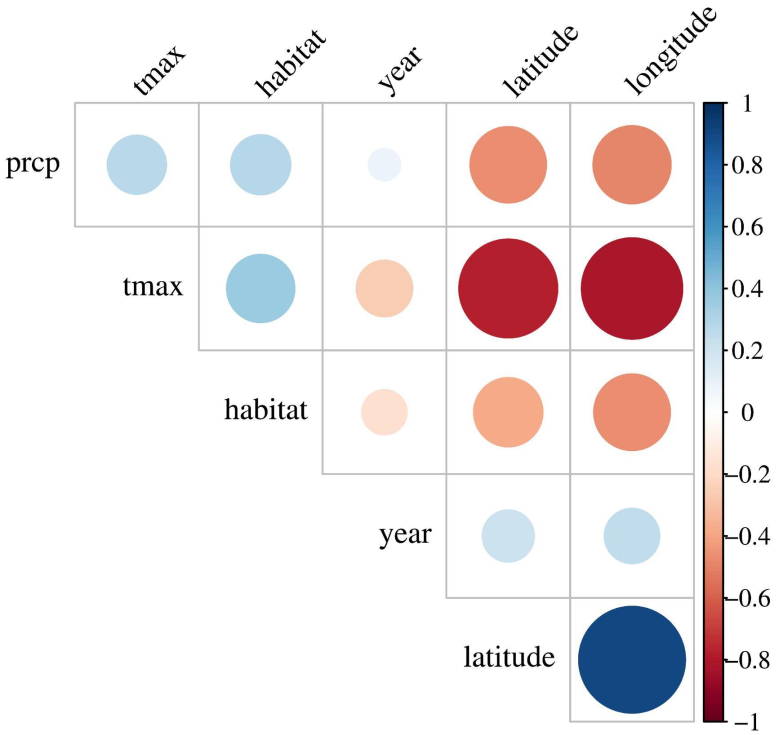

**Fig. S19.** Correlation matrix of the covariates from the multivariate model on haplotype change over time, space, habitat and climate (Fig. 5 analyses). All correlations were significant (Pearson's correlation test, *p*-value < 0.05). prcp-value = total precipitation for summer months, tmax = maximum temperature for summer months. The "habitat" covariate was turned to "1" for Agricultural and "0" for Natural habitats.

*Supplementary Tables*

**Table S1.** Multivariate models testing the effects of longitude, latitude and environment on drought resistance (**Figure 2C&D**).

| Model with latitude | Estimate | Std. Error | t | p-value |
|---|---|---|---|---|
| (Intercept) | 7.67E+00 | 23.227 | 0.33 | 0.7446 |
| Habitat | -9.25E-01 | 0.5283 | -1.751 | 0.0952 |
| Longitude | -1.53E-01 | 0.113 | -1.357 | 0.1899 |
| Latitude | -2.50E-01 | 0.3681 | -0.679 | 0.5047 |
| AIC | 8.61E+01 | | | |
| Model without latitude | Estimate | Std. Error | t | p-value |
| (Intercept) | -7.44E+00 | 6.59162 | -1.129 | 0.27172 |
| Longitude | -2.11E-01 | 0.07361 | -2.867 | 0.00923 |
| Habitat | -9.25E-01 | 0.52145 | -1.774 | 9.05E-02 |
| AIC | 8.47E+01 | | | |

**Table S2**. Results from the univariate models testing for trade-offs/positive associations between growth, herbicide resistance and drought tolerance. Plots are shown in Fig. S4.

| variable 1 | variable 2 | Experimental treatment | scale | slope (beta) | $R^2$ | *p*-value |
|---|---|---|---|---|---|---|
| growth | herbicide resistance | control | individual | 0.013 | 0.00013 | 0.8615 |
| growth | herbicide resistance | drought | individual | -0.078 | 0.0048 | 0.302 |
| growth | herbicide resistance | control | population | -0.294 | 0.14 | 0.087 |
| growth | herbicide resistance | drought | population | -0.172 | 0.045 | 0.317 |
| **drought resistance** | **herbicide resistance** | **drought** | **individual** | **0.50** | **0.025** | **0.0087** |
| **drought resistance** | **herbicide resistance** | **drought** | **population** | **1.50** | **0.36** | **0.0077** |
| growth | drought resistance | control | individual | 0.230 | 0.00713 | 0.19 |
| **growth** | **drought resistance** | **drought** | **individual** | **0.744** | **0.107** | **0.0000005549** |
| **growth** | **drought resistance** | **control** | **population** | **-2.14** | **0.3526** | **0.00357** |
| growth | drought resistance | drought | population | -0.212 | 0.03993 | 0.73 |

**Table S3**. Results from the individual multivariate mixed models testing for trade-offs/positive associations between growth, herbicide resistance and drought tolerance. Plots are shown in **Fig. S4**.

| **Random effects** | Name | Variance | Std.Dev. | | |
|---|---|---|---|---|---|
| Tray:Bench | (Intercept) | 0.02774 | 0.1666 | | |
| Bench | (Intercept) | 0.08008 | 0.283 | | |
| Residual | 6.06E+00 | 2.461 | | | |
| **Fixed effects** | Estimate | SE | DF | t | p-value |
| (Intercept) | 3.90317 | 6.39011 | 213.6207 | 0.611 | 0.542 |
| EPSPS | 0.18055 | 0.1748 | 212.14424 | 1.033 | 0.303 |
| Growth rate | 0.76683 | 0.14349 | 209.49625 | 5.344 | 2.36E-07 |
| Longitude | -0.05658 | 0.07621 | 212.62611 | -0.742 | 0.459 |
| Habitat | -1.03599 | 0.79421 | 214.61198 | -1.304 | 0.193 |
| Ancestry | 1.78E-01 | 1.04E+00 | 215.64492 | 0.172 | 0.864 |
| Habitat:Ancestry | 1.30E+00 | 1.02E+00 | 213.14108 | 1.275 | 0.204 |

**Table S4**. Response to drought depends on ancestry, regardless of ADMIXTURE or PC based metric in multivariate models. Models were implemented with two different hierarchical summaries of ancestry, at the individual and population levels.

| | **Ancestry** | Sum Sq | Df | F | value |
|---|---|---|---|---|---|
| | (Intercept) | 2.648 | 1 | 2.696 | 0.12011 |
| | longitude | 6.1124 | 1 | 6.2232 | 0.02393 |
| | latitude | 0.7919 | 1 | 0.8062 | 0.38255 |
| | ADMIXTURE proportion | 5.8556 | 1 | 5.9618 | 0.02661 |
| | env | 0.3516 | 1 | 0.358 | 0.55801 |
| **Population-level** | longitude:ADMIXTURE proportion | 7.0237 | 1 | 7.151 | 0.01663 |
| | Residuals | 15.715 | 16 | | |
| | **PC1** | | | | |
| | (Intercept) | 1.1793 | 1 | 1.2296 | 0.28387 |
| | meanPC1 | 6.8456 | 1 | 7.1379 | 0.01671 |
| | longitude | 2.8544 | 1 | 2.9763 | 0.10375 |
| | latitude | 0.4412 | 1 | 0.46 | 0.50731 |
| | env | 0.4217 | 1 | 0.4398 | 0.51668 |

| | | | | | |
|---|---|---|---|---|---|
| | meanPC1:longitude | 8.0677 | 1 | 8.4122 | 0.01043 |
| | Residuals | 15.3446 | 16 | | |
| Individual-level | **Ancestry** | | | | |
| | latitude | 5.1 | 1 | 0.5461 | 4.61E-01 |
| | longitude | 5.4 | 1 | 0.578 | 4.48E-01 |
| | env | 11.64 | 1 | 1.2462 | 2.65E-01 |
| | ADMIXTURE proportion | 53.27 | 1 | 5.7049 | 0.01759 |
| | longitude:ADMIXTURE proportion | 42.6 | 1 | 4.5614 | 3.36E-02 |
| | Residuals | 2558.68 | 274 | | |
| | **PC1** | | | | |
| | latitude | 1.98 | 1 | 0.2116 | 6.46E-01 |
| | longitude | 5.2 | 1 | 0.5543 | 0.4572 |
| | env | 12.8 | 1 | 1.365 | 0.24368 |
| | PC1 | 37.41 | 1 | 3.9895 | 4.68E-02 |
| | longitude:PC1 | 48.06 | 1 | 5.1252 | 0.02436 |
| | Residuals | 2569.08 | 274 | | |

**Table S5.** Variant type (as identified by snpEff) and potential functional effect of significant admixture mapping hits with drought tolerance.

| chromosome | pos | ref | alt | type | protein_&_species |
|---|---|---|---|---|---|
| 2 | 51130514 | A | T | missense_variant | ABCA2: ABC transporter A family member 2 (Arabidopsis thaliana); |
| 4 | 5060424 | A | T | missense_variant | HAT14: Homeobox-leucine zipper protein HAT14 (Arabidopsis thaliana) |
| 4 | 5075021 | A | T | missense_variant | Similar to TBR: Protein trichome birefringence (Arabidopsis thaliana) |
| 4 | 5075469 | A | T | missense_variant | Similar to TBR: Protein trichome birefringence (Arabidopsis thaliana) |
| 4 | 5075662 | A | T | splice_region_variant&intron_variant | Note=Similar to TBR: Protein trichome birefringence (Arabidopsis thaliana); |
| 5 | 37694747 | A | T | missense_variant | Similar to TDC2: Tryptophan decarboxylase TDC2 (Camptotheca acuminata) |
| 5 | 37694833 | A | T | missense_variant | Similar to TDC2: Tryptophan decarboxylase TDC2 (Camptotheca acuminata) |
| 5 | 37694869 | A | T | missense_variant | Similar to TDC2: Tryptophan decarboxylase TDC2 (Camptotheca acuminata) |
| 5 | 37694930 | A | T | missense_variant | Similar to TDC2: Tryptophan decarboxylase TDC2 (Camptotheca acuminata) |
| 5 | 37694938 | A | T | missense_variant | Similar to TDC2: Tryptophan decarboxylase TDC2 (Camptotheca acuminata) |
| 5 | 37695440 | A | T | missense_variant | Similar to TDC2: Tryptophan decarboxylase TDC2 (Camptotheca acuminata) |
| 5 | 37695515 | A | T | missense_variant | Similar to TDC2: Tryptophan decarboxylase TDC2 (Camptotheca acuminata) |
| 6 | 2332437 | A | T | missense_variant | Protein of unknown function; |
| 6 | 2332891 | A | T | missense_variant | Protein of unknown function; |
| 6 | 2333185 | A | T | missense_variant | Protein of unknown function; |
| 6 | 2396601 | A | T | missense_variant | Similar to At4g34215: Probable carbohydrate esterase At4g34215 (Arabidopsis thaliana); |
| 6 | 3055974 | A | T | missense_variant | IPR012951 Berberine/berberine-like |
| 6 | 3056020 | A | T | missense_variant | IPR012951 Berberine/berberine-like |
| 6 | 3458305 | A | T | missense_variant | Protein of unknown function |
| 7 | 6568735 | A | T | missense_variant | Protein of unknown function |
| 7 | 6571047 | A | T | missense_variant | Protein of unknown function |
| 7 | 6572134 | A | T | missense_variant | Protein of unknown function |
| 7 | 6572212 | A | T | missense_variant | Protein of unknown function |
| 7 | 6572213 | A | T | missense_variant | Protein of unknown function |
| 7 | 6572273 | A | T | missense_variant | Protein of unknown function |
| 7 | 6591276 | A | T | missense_variant | Protein of unknown function |
| 7 | 8207600 | A | T | missense_variant | Protein of unknown function |
| 7 | 8207667 | A | T | missense_variant | Protein of unknown function |
| 7 | 6568883 | A | T | missense_variant&splice_region_variant | Protein of unknown function; |
| 7 | 6572307 | A | T | stop_gained | Protein of unknown function; |
| 9 | 20981908 | A | T | missense_variant | Similar to RKD5: Protein RKD5 (Arabidopsis thaliana); |
| 9 | 20982118 | A | T | missense_variant | Similar to RKD5: Protein RKD5 (Arabidopsis thaliana); |
| 9 | 20981232 | A | T | stop_gained | Similar to RKD5: Protein RKD5 (Arabidopsis thaliana); |
| 9 | 20981616 | A | T | stop_gained | Similar to RKD5: Protein RKD5 (Arabidopsis thaliana); |
| 10 | 22292099 | A | T | missense_variant | Similar to NOP2B: 26S rRNA (cytosine-C(5))-methyltransferase NOP2B (Arabidopsis thaliana); |
| 10 | 22292115 | A | T | missense_variant | Similar to NOP2B: 26S rRNA (cytosine-C(5))-methyltransferase NOP2B (Arabidopsis thaliana); |
| 10 | 22292290 | A | T | missense_variant | Similar to NOP2B: 26S rRNA (cytosine-C(5))-methyltransferase NOP2B (Arabidopsis thaliana); |
| 10 | 22301287 | A | T | stop_gained | Similar to PER17: Peroxidase 17 (Arabidopsis thaliana); |
| 13 | 20668702 | A | T | missense_variant | Protein of unknown function |
| 13 | 20667144 | A | T | missense_variant&splice_region_variant | Protein of unknown function; |

| | | | | | |
|---|---|---|---|---|---|
| 13 | 20666046 | A | T | splice_region_variant&intron_variant | Protein of unknown function; |
| 13 | 20667807 | A | T | splice_region_variant&intron_variant | Protein of unknown function; |
| 16 | 22780805 | A | T | missense_variant | Similar to APSR1: Protein ALTERED PHOSPHATE STARVATION RESPONSE 1 (Arabidopsis thaliana); |
| 16 | 22781009 | A | T | missense_variant | Similar to APSR1: Protein ALTERED PHOSPHATE STARVATION RESPONSE 1 (Arabidopsis thaliana); |
| 16 | 22782693 | A | T | missense_variant | Similar to APSR1: Protein ALTERED PHOSPHATE STARVATION RESPONSE 1 (Arabidopsis thaliana); |
| 16 | 22783372 | A | T | missense_variant | Similar to APSR1: Protein ALTERED PHOSPHATE STARVATION RESPONSE 1 (Arabidopsis thaliana); |
| 16 | 22783480 | A | T | missense_variant | Similar to APSR1: Protein ALTERED PHOSPHATE STARVATION RESPONSE 1 (Arabidopsis thaliana); |
| 16 | 22784473 | A | T | missense_variant | Similar to APSR1: Protein ALTERED PHOSPHATE STARVATION RESPONSE 1 (Arabidopsis thaliana); |
| 16 | 22784515 | A | T | missense_variant | Similar to APSR1: Protein ALTERED PHOSPHATE STARVATION RESPONSE 1 (Arabidopsis thaliana); |
| 16 | 22784532 | A | T | missense_variant | Atub_193_hap2_00024293-RA:cds |
| 16 | 22801329 | A | T | missense_variant | Similar to DDB1: DNA damage-binding protein 1 (Solanum lycopersicum); |
| 16 | 22802704 | A | T | missense_variant | Similar to DDB1: DNA damage-binding protein 1 (Solanum lycopersicum); |
| 16 | 22808757 | A | T | missense_variant | Similar to DDB1: DNA damage-binding protein 1 (Solanum lycopersicum); |
| 16 | 22808916 | A | T | missense_variant | Similar to DDB1: DNA damage-binding protein 1 (Solanum lycopersicum); |
| 16 | 22813743 | A | T | missense_variant | Similar to DDB1: DNA damage-binding protein 1 (Solanum lycopersicum); |
| 16 | 22815726 | A | T | missense_variant | Similar to DDB1: DNA damage-binding protein 1 (Solanum lycopersicum); |
| 16 | 22820109 | A | T | missense_variant | Similar to DDB1: DNA damage-binding protein 1 (Solanum lycopersicum); |
| 16 | 22818417 | A | T | stop_gained | Similar to DDB1: DNA damage-binding protein 1 (Solanum lycopersicum); |
| 16 | 22820133 | A | T | stop_gained | Similar to DDB1: DNA damage-binding protein 1 (Solanum lycopersicum); |
| 16 | 22829741 | A | T | missense_variant | Similar to ABCC1: ABC transporter C family member 1 (Arabidopsis thaliana); |
| 16 | 22830323 | A | T | missense_variant | Similar to ABCC1: ABC transporter C family member 1 (Arabidopsis thaliana); |
| 16 | 22830452 | A | T | missense_variant | Similar to ABCC1: ABC transporter C family member 1 (Arabidopsis thaliana); |
| 16 | 22830466 | A | T | missense_variant | Similar to ABCC1: ABC transporter C family member 1 (Arabidopsis thaliana); |
| 16 | 22831915 | A | T | missense_variant | Similar to ABCC1: ABC transporter C family member 1 (Arabidopsis thaliana); |
| 16 | 22942706 | A | T | missense_variant | Similar to REM4.1: Remorin 4.1 (Oryza sativa subsp. japonica); |
| 16 | 22942714 | A | T | missense_variant | Similar to REM4.1: Remorin 4.1 (Oryza sativa subsp. japonica); |
| 16 | 22942822 | A | T | missense_variant | Similar to REM4.1: Remorin 4.1 (Oryza sativa subsp. japonica); |
| 16 | 22943115 | A | T | missense_variant | Similar to REM4.1: Remorin 4.1 (Oryza sativa subsp. japonica); |
| 16 | 22955554 | A | T | missense_variant | Similar to CIPK23: CBL-interacting serine/threonine-protein kinase 23 (Arabidopsis thaliana); |
| 16 | 22955666 | A | T | missense_variant | Similar to CIPK23: CBL-interacting serine/threonine-protein kinase 23 (Arabidopsis thaliana); |
| 16 | 22955727 | A | T | missense_variant | Similar to CIPK23: CBL-interacting serine/threonine-protein kinase 23 (Arabidopsis thaliana); |
| 16 | 22999020 | A | T | missense_variant | Similar to TCP2: Transcription factor TCP2 (Arabidopsis thaliana); |
| 16 | 22999363 | A | T | missense_variant | Similar to TCP2: Transcription factor TCP2 (Arabidopsis thaliana); |
| 16 | 22841981 | A | T | splice_region_variant&intron_variant | Note=Similar to ABCC1: ABC transporter C family member 1 (Arabidopsis thaliana); |

| | | | | | |
|---|---|---|---|---|---|
| 16 | 22879935 | A | T | splice_region_variant&intron_variant | Protein of unknown function; |
| 16 | 22941059 | A | T | splice_region_variant&intron_variant | Similar to REM4.1: Remorin 4.1 (Oryza sativa subsp. japonica); |
| 16 | 22939421 | A | T | stop_gained | Similar to REM4.1: Remorin 4.1 (Oryza sativa subsp. japonica); |
| 16 | 22955293 | A | T | splice_region_variant&intron_variant | Similar to CIPK23: CBL-interacting serine/threonine-protein kinase 23 (Arabidopsis thaliana); |
| 16 | 22889639 | A | T | splice_region_variant&synonymous_variant | Similar to POPTRDRAFT_798217: CASP-like protein 1B2 (Populus trichocarpa); |
| 16 | 22890784 | A | T | stop_gained | Similar to POPTRDRAFT_798217: CASP-like protein 1B2 (Populus trichocarpa); |
| 16 | 22886861 | A | T | stop_gained | Protein of unknown function; |

**Table S6.** Biological processes inferred from gene ontology analysis of  893 admixture mapping hits for drought-tolerance.

| GO biological process complete | Arabidopsis thaliana - REFLIST | Fold Enrichment | raw *p*-value |
|---|---|---|---|
| mRNA 3'-end processing by stem-loop binding and cleavage (GO:0006398) | 2 | > 100 | 2.26E-03 |
| (+)-abscisic acid D-glucopyranosyl ester transmembrane transport (GO:1902418) | 2 | > 100 | 2.26E-03 |
| phytochromobilin biosynthetic process (GO:0010024) | 3 | > 100 | 3.38E-03 |
| phytochromobilin metabolic process (GO:0051202) | 3 | > 100 | 3.38E-03 |
| phenylacetate catabolic process (GO:0010124) | 3 | > 100 | 3.38E-03 |
| glucoside transport (GO:0042946) | 3 | > 100 | 3.38E-03 |
| xenobiotic catabolic process (GO:0042178) | 4 | > 100 | 4.51E-03 |
| xenobiotic metabolic process (GO:0006805) | 5 | > 100 | 5.63E-03 |
| cellular response to xenobiotic stimulus (GO:0071466) | 5 | > 100 | 5.63E-03 |
| histone mRNA metabolic process (GO:0008334) | 5 | > 100 | 5.63E-03 |
| glycoside transport (GO:1901656) | 5 | > 100 | 5.63E-03 |
| xenobiotic transmembrane transport (GO:0006855) | 6 | > 100 | 6.75E-03 |
| riboflavin biosynthetic process (GO:0009231) | 8 | > 100 | 8.99E-03 |
| arsenite transport (GO:0015700) | 8 | > 100 | 8.99E-03 |
| positive regulation of photomorphogenesis (GO:2000306) | 8 | > 100 | 8.99E-03 |
| positive regulation of development, heterochronic (GO:0045962) | 8 | > 100 | 8.99E-03 |
| potassium ion import across plasma membrane (GO:1990573) | 8 | > 100 | 8.99E-03 |
| chloroplast-nucleus signaling pathway (GO:0010019) | 9 | 98.48 | 1.01E-02 |
| response to xenobiotic stimulus (GO:0009410) | 9 | 98.48 | 1.01E-02 |
| riboflavin metabolic process (GO:0006771) | 9 | 98.48 | 1.01E-02 |
| pollen tube reception (GO:0010483) | 11 | 80.57 | 1.23E-02 |
| flavin-containing compound biosynthetic process (GO:0042727) | 12 | 73.86 | 1.35E-02 |
| biological process involved in intraspecies interaction between organisms (GO:0051703) | 14 | 63.31 | 1.57E-02 |
| intracellular manganese ion homeostasis (GO:0030026) | 14 | 63.31 | 1.57E-02 |
| flavin-containing compound metabolic process (GO:0042726) | 15 | 59.09 | 1.68E-02 |
| mRNA 3'-end processing (GO:0031124) | 15 | 59.09 | 1.68E-02 |
| regulation of morphogenesis of a branching structure (GO:0060688) | 15 | 59.09 | 1.68E-02 |
| regulation of secondary shoot formation (GO:2000032) | 15 | 59.09 | 1.68E-02 |
| manganese ion homeostasis (GO:0055071) | 16 | 55.39 | 1.79E-02 |
| response to nutrient (GO:0007584) | 16 | 55.39 | 1.79E-02 |
| regulation of plant organ formation (GO:1905428) | 19 | 46.65 | 2.12E-02 |
| response to arsenic-containing substance (GO:0046685) | 19 | 46.65 | 2.12E-02 |
| rRNA base methylation (GO:0070475) | 19 | 46.65 | 2.12E-02 |
| inorganic cation import across plasma membrane (GO:0098659) | 19 | 46.65 | 2.12E-02 |
| inorganic ion import across plasma membrane (GO:0099587) | 20 | 44.31 | 2.23E-02 |
| root hair initiation (GO:0048766) | 20 | 44.31 | 2.23E-02 |
| snRNA processing (GO:0016180) | 21 | 42.2 | 2.34E-02 |
| chloroplast accumulation movement (GO:0009904) | 21 | 42.2 | 2.34E-02 |
| potassium ion transport (GO:0006813) | 51 | 34.76 | 1.52E-03 |
| snRNA metabolic process (GO:0016073) | 27 | 32.83 | 3.00E-02 |
| dicarboxylic acid biosynthetic process (GO:0043650) | 28 | 31.65 | 3.11E-02 |
| xenobiotic transport (GO:0042908) | 57 | 31.1 | 1.89E-03 |
| rRNA methylation (GO:0031167) | 29 | 30.56 | 3.22E-02 |
| establishment of plastid localization (GO:0051667) | 29 | 30.56 | 3.22E-02 |
| chloroplast relocation (GO:0009902) | 29 | 30.56 | 3.22E-02 |
| import across plasma membrane (GO:0098739) | 33 | 26.86 | 3.66E-02 |
| potassium ion transmembrane transport (GO:0071805) | 34 | 26.07 | 3.77E-02 |
| chloroplast localization (GO:0019750) | 34 | 26.07 | 3.77E-02 |
| maturation of LSU-rRNA (GO:0000470) | 34 | 26.07 | 3.77E-02 |
| plastid localization (GO:0051644) | 35 | 25.32 | 3.88E-02 |
| fatty acid beta-oxidation (GO:0006635) | 36 | 24.62 | 3.99E-02 |
| positive gravitropism (GO:0009958) | 36 | 24.62 | 3.99E-02 |
| regulation of photomorphogenesis (GO:0010099) | 40 | 22.16 | 4.42E-02 |
| fatty acid oxidation (GO:0019395) | 40 | 22.16 | 4.42E-02 |

| | | | |
|---|---|---|---|
| lignin biosynthetic process (GO:0009809) | 41 | 21.62 | 4.53E-02 |
| rRNA modification (GO:0000154) | 43 | 20.61 | 4.74E-02 |
| fatty acid catabolic process (GO:0009062) | 43 | 20.61 | 4.74E-02 |
| inorganic cation transmembrane transport (GO:0098662) | 204 | 8.69 | 2.21E-02 |
| metal ion transport (GO:0030001) | 227 | 7.81 | 2.70E-02 |
| monoatomic cation homeostasis (GO:0055080) | 242 | 7.32 | 3.04E-02 |
| inorganic ion transmembrane transport (GO:0098660) | 256 | 6.92 | 3.37E-02 |
| monoatomic ion homeostasis (GO:0050801) | 260 | 6.82 | 3.46E-02 |
| monoatomic cation transport (GO:0006812) | 264 | 6.71 | 3.56E-02 |
| monoatomic ion transport (GO:0006811) | 293 | 6.05 | 4.30E-02 |
| phosphorylation (GO:0016310) | 528 | 5.04 | 2.13E-02 |
| establishment of localization (GO:0051234) | 2263 | 3.13 | 2.95E-03 |
| localization (GO:0051179) | 2434 | 2.91 | 4.63E-03 |
| transport (GO:0006810) | 2137 | 2.9 | 8.52E-03 |

**Table S7.** ANOVA results from multivariate logistic and linear models of joint allele frequency change of all drought loci across sequenced herbarium samples spanning spatiotemporal scales, before and after accounting for collection habitat (classified as disturbed, agricultural, or natural) and its interaction with time. Coefficients from the model shown both as the raw value and as the homozygous alternate fitness advantage (2 x β = $B_{AA}$ or $s_{AA}$).

| | Linear model | β | $\beta_{AA}$ | Sums of Squares | DF | F | *p*-value |
|---|---|---|---|---|---|---|---|
| No habitat | (Intercept) | 23.5 | - | 0.935 | 1 | 7.4246 | 0.0065371 |
| | tmax | -0.741 | -1.49 | 1.059 | 1 | 8.4072 | 0.0038124 |
| | year | -0.01335 | -2.67E-02 | 1.126 | 1 | 8.941 | 0.0028512 |
| | prcp | 0.007795 | 1.56E-02 | 0.381 | 1 | 3.0282 | 0.0821114 |
| | locus | - | - | 6.421 | 32 | 1.5935 | 0.0198677 |
| | samp_lat | 0.04234 | 4.23E-02 | 1.745 | 1 | 13.8551 | 0.0002076 |
| | samp_lon | -0.00966 | -1.93E-02 | 0.496 | 1 | 3.9362 | 0.0475087 |
| | tmax:year | 0.0003935 | 7.87E-04 | 1.096 | 1 | 8.7026 | 0.0032455 |
| | year:prcp | -0.000003979 | -7.96E-06 | 0.375 | 1 | 2.9817 | 0.0844971 |
| | Residuals | | | 136.996 | 1088 | | |
| With habitat | Linear model | β | $\beta_{AA}$ | Sums of Squares | DF | F | *p*-value |
| | (Intercept) | -1.82E+01 | - | 0.187 | 1 | 1.5924 | 2.07E-01 |
| | tmax | 2.64E-01 | 5.28E-01 | 0.06 | 1 | 0.5108 | 4.75E-01 |
| | year | 7.09E-03 | 1.42E-02 | 0.108 | 1 | 0.9221 | 3.37E-01 |
| | prcp | 1.15E-02 | 2.30E-02 | 0.526 | 1 | 4.4895 | 3.43E-02 |
| | env | - | - | 0.697 | 2 | 2.9705 | 5.17E-02 |
| | locus | - | - | 5.524 | 32 | 1.4721 | 4.49E-02 |
| | latitude | 6.42E-02 | 1.28E-01 | 3.532 | 1 | 30.1238 | 5.14E-08 |
| | longitude | -1.45E-02 | -2.90E-02 | 0.952 | 1 | 8.1198 | 4.47E-03 |
| | tmax:year | -1.19E-04 | -2.38E-04 | 0.046 | 1 | 0.3964 | 5.29E-01 |
| | year:prcp | -5.81E-06 | -1.16E-05 | 0.51 | 1 | 4.3487 | 3.73E-02 |
| | year:env | - | - | 0.7 | 2 | 2.9838 | 5.11E-02 |
| | Residuals | | | 117.488 | 1002 | | |

| | Logistic model | β | $s_{AA}$ | Chisq | DF | *p*-value |
|---|---|---|---|---|---|---|
| No habitat | tmax | 9.88E+01 | 1.98E+02 | 4.5502 | 1 | 0.032915 |
| | year | -3.22E+00 | -6.44E+00 | 4.8359 | 1 | 0.027872 |
| | prcp | -5.75E-02 | -1.15E-01 | 1.6811 | 1 | 0.194773 |
| | locus | 3.42E-02 | 6.84E-02 | 27.236 | 32 | 0.706567 |
| | latitude | 1.83E-01 | 3.66E-01 | 7.5157 | 1 | 0.006116 |
| | longitude | -4.12E-02 | -8.24E-02 | 2.0563 | 1 | 0.151575 |
| | tmax:year | 1.70E-03 | 3.40E-03 | 4.7095 | 1 | 0.029996 |
| | year:prcp | -1.74E-05 | -3.48E-05 | 1.6545 | 1 | 0.198346 |
| | Logistic model | β | $s_{AA}$ | Chisq | DF | *p*-value |
| With habitat | tmax | -8.06E+01 | -1.61E+02 | 0.2481 | 1 | 0.61843 |
| | year | 1.12E+00 | 2.24E+00 | 0.4621 | 1 | 0.49664 |
| | prcp | 3.05E-02 | 6.10E-02 | 2.4166 | 1 | 0.12006 |
| | locus | 5.12E-02 | 1.02E-01 | 23.3951 | 32 | 0.86547 |
| | latitude | 2.78E-01 | 5.56E-01 | 15.277 | 1 | 9.28E-05 |
| | longitude | -6.28E-02 | -1.26E-01 | 4.1616 | 1 | 0.04135 |
| | env | - | - | 2.9956 | 2 | 0.22363 |
| | tmax:year | -5.04E-04 | -1.01E-03 | 0.1915 | 1 | 0.6617 |
| | year:prcp | -2.59E-05 | -5.18E-05 | 2.3425 | 1 | 0.12589 |
| | year:env | - | - | 3.0081 | 2 | 0.22223 |
